# Cell-type-specific chromatin occupancy by the pioneer factor Zelda drives key developmental transitions in *Drosophila*

**DOI:** 10.1101/2021.03.10.434844

**Authors:** Elizabeth D. Larson, Hideyuki Komori, Tyler J. Gibson, Cyrina M. Ostgaard, Danielle C. Hamm, Jack M. Schnell, Cheng-Yu Lee, Melissa M. Harrison

## Abstract

During *Drosophila* embryogenesis, the essential pioneer factor Zelda defines hundreds of *cis*-regulatory regions and in doing so reprograms the zygotic transcriptome. While Zelda is essential later in development, it is unclear how the ability of Zelda to define *cis*-regulatory regions is shaped by cell-type-specific chromatin architecture. Asymmetric division of neural stem cells (neuroblasts) in the fly brain provide an excellent paradigm for investigating the cell-type-specific functions of this pioneer factor. We show that Zelda synergistically functions with Notch to maintain neuroblasts in an undifferentiated state. Zelda misexpression reprograms progenitor cells to neuroblasts, but this capacity is limited by transcriptional repressors critical for progenitor commitment. Zelda genomic occupancy in neuroblasts is reorganized as compared to the embryo, and this reorganization is driven by differences in chromatin accessibility and cofactor availability. We propose that Zelda regulates essential transitions in the neuroblasts and embryo through a shared gene-regulatory network by defining cell-type-specific enhancers.

## Introduction

During development, the genome is differentially interpreted to give rise to thousands of distinctive cell types. Once terminally differentiated, cells within an organism are generally incapable of transitioning to a less differentiated fate. By contrast, in cell culture the addition of a cocktail of transcription factors can reprogram differentiated cells back to a pluripotent state. Many of these reprogramming factors, including Oct4, Sox2 and Klf4, function as pioneer transcription factors: a specialized set of transcription factors that can bind DNA within the context of nucleosomes, facilitate chromatin accessibility and subsequent binding by additional transcription factors (Iwafuchi-Doi, 2019; Iwafuchi-Doi and Zaret, 2014; Soufi et al., 2012, 2015; Zaret and Carroll, 2011). These features of reprogramming pioneer factors allow them to gain access to silenced regions of the genome and drive new gene expression profiles to change cell fate. Indeed, misexpression of pioneer factors within an organism leads to dramatic gene expression changes that can cause disease (Ben-Porath et al., 2008; Geng et al., 2012; Tsai et al., 2011; Wang and Herlyn, 2015; Young et al., 2013). Despite the ability of these factors to engage silenced portions of the genome, there are barriers to pioneer-factor binding and efficient reprogramming (Chronis et al., 2017; Donaghey et al., 2018; Larson et al., 2021; Liu and Kraus, 2017; Mayran et al., 2018; Soufi et al., 2012; Swinstead et al., 2016; Zaret and Mango, 2016). Many studies have leveraged the advantages of cell culture systems to identify impediments to pioneer-factor binding and reprogramming. However, many fewer studies have identified limitations to pioneer factor-driven cell fate changes within the context of an entire, developing organism.

Immediately following fertilization, the specified germ cells must be reprogrammed to form the totipotent cells that can ultimately differentiate to generate a new organism. During this time, the zygotic genome is transcriptionally silent, and maternally deposited mRNAs and proteins control early embryonic development (Laver et al., 2015; Schulz and Harrison, 2019; Vastenhouw et al., 2019; Yartseva and Giraldez, 2015). This maternal-to-zygotic transition (MZT) is necessary for development, and is orchestrated, in part, by factors that reprogram the zygotic genome for transcriptional activation (Schulz and Harrison, 2019; Vastenhouw et al., 2019). Factors that activate the zygotic genome have been identified in many species and all share essential features of pioneer factors. Perhaps the best characterized of these transcriptional regulators is Zelda (Zld), which we and others have shown functions as a pioneer factor to reprogram the early embryonic genome in *Drosophila melanogaster* (Foo et al., 2015; Hamm et al., 2015, 2017; Harrison et al., 2011; Liang et al., 2008; Nien et al., 2011; Schulz et al., 2015).

In the early embryo, Zld binding is driven strongly by sequence with between 40-65% of the canonical Zld-binding motifs (CAGGTAG) bound by Zld during the MZT (Harrison et al., 2011). This binding is distinctive even for pioneer transcription factors (Donaghey et al., 2018; Hurtado et al., 2011; Lupien et al., 2008; Tsankov et al., 2015). For example, the extensively studied pioneer factor FOXA2 only binds ~10% of its motifs in a variety of different cell types (Donaghey et al., 2018). The chromatin environment in the early embryo is naïve as compared to later in development (Hamm and Harrison, 2018), and this may contribute to the widespread occupancy of CAGGTAG motifs by Zld at this stage of development. In the *Drosophila* embryo, similar to other organisms, there are relatively few post-translational modifications to the histone proteins, and the genome is packaged by a unique linker histone dBigH1, which is essential for proper development (Li et al., 2014; Pérez-Montero et al., 2013; Schulz and Harrison, 2019). In addition, early development is characterized by a series of 13, rapid, semi-synchronous nuclear divisions that each occur over approximately 10 minutes and are comprised of only synthesis (S) and division (M) phases (Hamm and Harrison, 2018). While, it is possible that the reprogramming function of Zld requires these distinctive properties of early development, Zld is also necessary for development after the MZT (Liang et al., 2008). It remains unclear whether Zld defines *cis*-regulatory regions in tissues outside the early embryo and if so, how this activity is regulated by the cell-type specific chromatin established during development.

Asymmetric division of neural stem cells (neuroblasts) in the larval brain provide an excellent *in vivo* system for investigating temporal regulation of enhancer activity. In the larval brain lobe, there are predominantly two types of neural stem cell populations: type I and type II (Bello et al., 2008; Boone and Doe, 2008; Bowman et al., 2008). Both types of neuroblasts undergo asymmetric division to self-renew and to generate a descendant that exits the multi-potent state and begin to differentiate. While type I neuroblasts directly contribute to neurogenesis, type II neuroblasts divide asymmetrically to self-renew and to generate a sibling cell that commits to an intermediate neural progenitor (INP) identity and functions as a transit-amplifying cell. One of the newly born neuroblast progeny, marked by the absence of Deadpan (Dpn) expression, first transitions into a non-Asense-expressing (Dpn^−^Ase^−^) immature INP, and then, after three to four hours, to an Asense-expressing (Dpn^−^Ase^+^) immature INP. Dpn is re-expressed in mature INPs (Dpn^+^Ase+) that undergo 6-8 rounds of asymmetric division to exclusively generate differentiated cell types (Figure 1A). Thus, despite Dpn expression and the capacity to undergo a limited number of asymmetric divisions, INPs lack the functional characteristics of the type II neuroblast from which they are derived. These molecularly defined intermediate stages of INP commitment provide a powerful system to investigate the temporal control of enhancer activity as stem cells exit the undifferentiated state.

**Figure 1.**
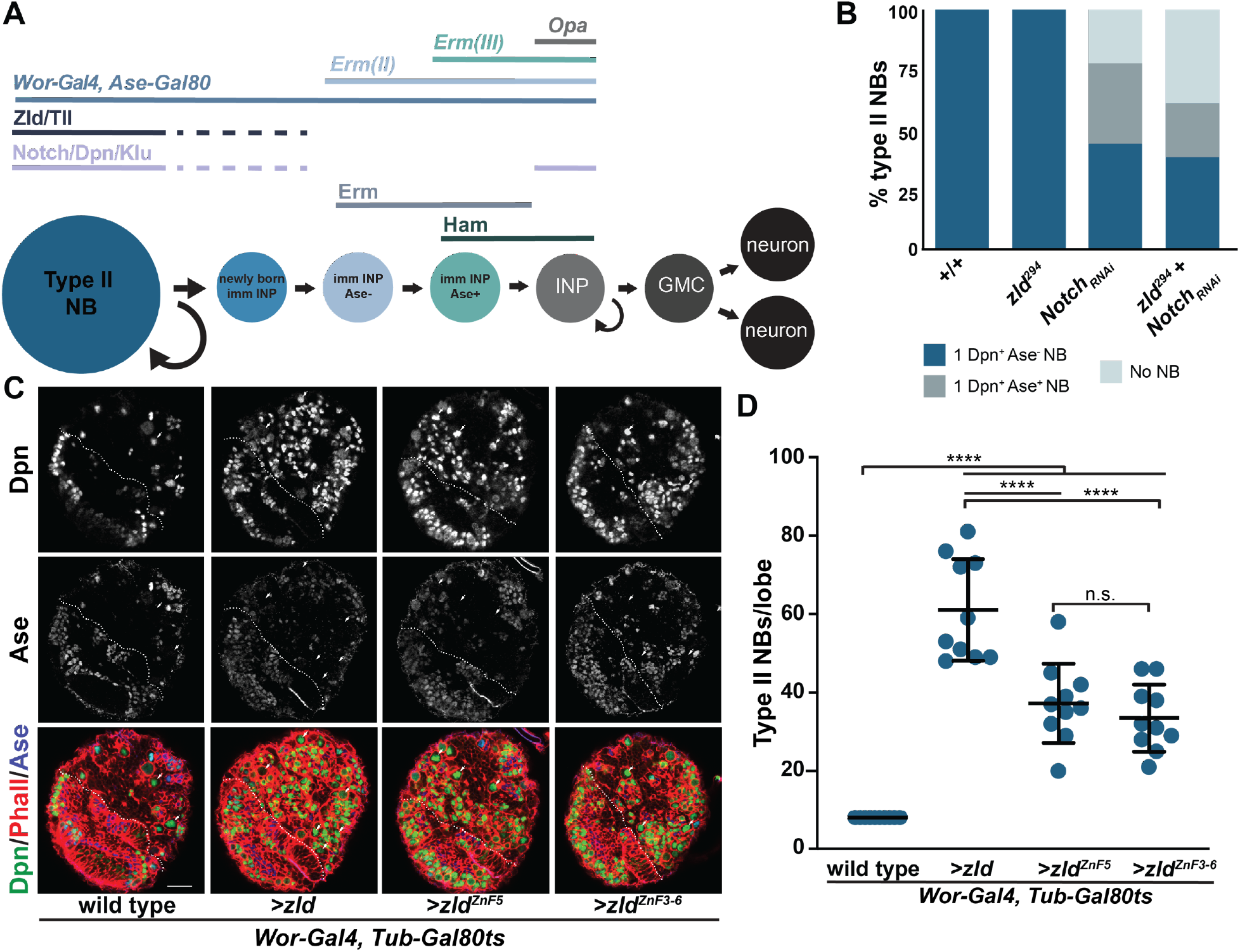
Zld promotes the undifferentiated type II neuroblast fate. A. Expression of self-renewal factors (Notch/Dpn/Klu), Tll, Zld, Erm and Ham and cell-type specific drivers along the type II neuroblast (NB) lineage. B. Percent of clones of the indicated genotype with neuroblasts (NB) expressing Dpn and/or Ase or lacking cells expressing these neuroblast markers. For +/+ n = 14 clones, *zld^294^* n = 16 clones, *Notch_RNAi_* n = 9 clones, *zld^294^* + *Notch_RNAi_* n = 13 clones. C. Immunostaining of third instar larval brain lobes without transgene expression (wild type), expression of ectopic Zld (>*zld*), expression of Zld with a mutation in the fifth zinc finger (>*zld^ZnF5^*), and expression of Zld with mutations in zinc fingers 3-6 (>*zld^Zn3-6^*) in type I and type II neuroblasts (*Wor-Gal4, Tub-Gal80ts*) at 33°C. White dashed line separates the optic lobe from the brain, and the white arrows indicate a selection of Dpn^+^, Ase^−^ type II neuroblasts. Scale bar, 20μm. D. Quantification of type II neuroblasts per lobe for the genotypes indicated. Wild-type (8 ± 0), >*zld* (61.1 ± 13), >*zld^ZnF5^* (37.3 ±10.1), and >*zld^Zn3-6^* (33.5 ± 8.5). Error bars show the standard deviation. For each genotype: n =10. Comparisons were performed with a one-way ANOVA with post hoc Tukey’s HSD test; n.s. = p-value > 0.05, **** = p-value ≤ 0.0001. See also Figure S1.

As in many stem cell populations, Notch signaling is essential for maintaining type II neuroblasts in an undifferentiated state: loss of Notch signaling causes premature commitment of neural stem cells to INP fate, and constitutive activation leads to extra neuroblasts (Janssens and Lee, 2014). However, despite reactivation of Notch signaling in the mature INPs, these cells do not revert back to stem cells. Downregulation of Notch signaling in newly born INPs allows the transcriptional repressors Earmuff (Erm) and Hamlet (Ham) to become sequentially activated during INP commitment (Eroglu et al., 2014; Janssens et al., 2014). Erm and Ham function through histone deacetylase 3 (Hdac3) to silence expression of *tailless (tll)*, which encodes a master regulator of type II neuroblast fate (Rives-Quinto et al., 2020). Suppressor of Hairless (Su(H)), the DNA-binding partner of Notch, binds the *tll* locus in larval brain neuroblasts, suggesting that *tll* is a Notch target (Zacharioudaki et al., 2016). Thus, in the INPs Erm- and Ham-mediated silencing of *tll* prevents reactivation of Notch signaling from inducing *tll* expression and driving their reversion to neuroblasts. Consistent with this model, INPs in *erm* or *ham*-mutant brains spontaneously revert to type II neuroblasts and knocking down Notch function can suppress the supernumerary neuroblast formation in these mutant brains (Rives-Quinto et al., 2020; Weng et al., 2010). These results support a model whereby sequential silencing by Erm and Ham during INP commitment renders the *tll* locus refractory to aberrant activation in INPs, but the precise mechanisms and the enhancers upon which these repressors act remains unclear.

Zld has been previously shown to be expressed in type II neuroblasts, but not in their progeny (Reichardt et al., 2018). Thus, the type II neuroblast lineage provides a powerful system to investigate the function of this pioneer factor in a tissue apart from the early embryo. We demonstrate that Zld functions with Notch signaling to maintain type II neuroblasts in an undifferentiated state. Exogenous expression of Zld during INP commitment can promote INP reversion into neuroblasts. However, changes to the chromatin state mediated by Erm and Ham limit this reprogramming capacity. We show that in type II neuroblasts chromatin occupancy by Zld is reorganized as compared to the embryo and that this reorganization is likely driven by chromatin accessibility. Nonetheless, target genes such as *dpn* and *tll* are shared between developmental stages. We propose that Zld drives key developmental transitions in the neuroblast lineage and the early embryo through a shared gene regulatory network by defining cell-type specific enhancers.

## Results

### Zelda functions synergistically with Notch to promote an undifferentiated state

Despite Zld expression being limited to the type II neuroblast, a previous study reported that knocking down *zld* function had no detectable effect on the neuroblast, but instead resulted in defects in INP proliferation (Reichardt et al., 2018). This discrepancy between the Zld expression pattern and the phenotype induced by RNAi-mediated *zld* knockdown prompted us to re-evaluate the role of *zld* in the larval brain (Figure 1A). We therefore assessed the identity of cells in mosaic clones derived from single *zld-*null mutant (*zld^294^*) neuroblasts using well-defined cell-fate markers. *zld*-mutant clones contained a single neuroblast and 15.4 ± 7 INPs, indistinguishable from wild-type clones (16.1 ± 6.2 INPs) (Figure 1B, Figure S1; for each genotype n = 10 clones). These data indicate that Zld is either dispensable in the type II neuroblast lineage or redundant pathways compensate for the absence of Zld.

Notch signaling plays an essential role in maintaining type II neuroblasts in an undifferentiated state (Xiao et al., 2012a; Zacharioudaki et al., 2012; Zhu et al., 2012). We therefore tested whether Zld might act synergistically with Notch signaling to regulate type II neuroblasts. For this purpose, we used a sensitized genetic background in which Notch function was reduced by RNAi. 22.2% of *Notch*-RNAi type II neuroblast clones lacked identifiable neuroblasts, and 33.3% of the clones contain neuroblasts with reduced cell diameter and Ase expression; two characteristics indicative of differentiation (n = 9 clones; Figure 1B). Simultaneous removal of *zld* enhanced this phenotype. 38.5% of *Notch*-RNAi, *zld*-mutant clones contained no neuroblasts, and 23% of clones contained neuroblasts with markers indicative of differentiation (n = 13 clones; Figure 1B). These data support a model in which Zld functions together with Notch to maintain type II neuroblasts in an undifferentiated state.

To further test if Zld promotes an undifferentiated state in type II neuroblasts, we overexpressed Zld using a series of UAS-*zld* transgenes under the control of a heat-inducible pan-neuroblast Gal4 driver (*Wor-Gal4, TubGal80^ts^*). While wild-type control brain lobes invariably contained 8 type II neuroblasts, 72 hours of Zld overexpression resulted in 61.1 ± 13 type II neuroblasts per lobe (n = 10 brains per genotype; Figure 1C,D). Zld is a zinc-finger transcription factor that binds to DNA in a sequence-specific manner that depends on a cluster of four zinc fingers in the C-terminus of the protein (Hamm et al., 2015). Mutation of either a single zinc finger or all four zinc fingers abrogates the ability of Zld to bind DNA and activate gene expression (Hamm et al., 2015). To determine if Zld overexpression drives this supernumerary type II neuroblast formation by binding DNA and regulating of gene expression, we overexpressed Zld protein with either a mutation in a single zinc finger of the DNA-binding domain (ZnF5) or all four zinc fingers of the DNA-binding domain mutated (ZnF3-6). Overexpressing either protein resulted in supernumerary type II neuroblast formation (37.3 ± 10.1 and 33.5 ± 8.5 type II neuroblasts per lobe, respectively; n = 10 brains per genotype), but expression of either mutant protein generated significantly fewer supernumerary neuroblasts than overexpression of wild-type Zld (Figure 1C,D). Together these data suggest that Zld promotes an undifferentiated state in type II neuroblasts and that this function is at least partially dependent on the ability of Zld to bind DNA in a sequence-specific manner.

### Zelda promotes an undifferentiated state in type II neuroblasts by activating *dpn* expression

Our data showed that Zld promotes an undifferentiated state in type II neuroblasts by functioning in parallel to or downstream of Notch. Aberrant activation of Notch signaling in either immature or mature INPs drives supernumerary neuroblast formation (Farnsworth et al., 2015). If Zld acts downstream of Notch to promote the undifferentiated state, then Zld misexpression in either immature INPs or mature INPs should similarly induce supernumerary neuroblasts. To test this, we induced Zld expression in different cell types along the neuroblast lineage and scored for supernumerary neuroblasts. Because no Gal4 drivers are exclusively active in immature INPs, we compared the effects of Zld misexpression in both immature INPs and mature INPs (driven by *Erm-Gal4*) to expression only in INPs (driven by *Opa-Gal4*) (Figure 1A). Zld overexpression throughout the type II neuroblast lineage driven by the *Wor-Gal4, Ase-Gal80* led to 44.5 ± 12.5 type II neuroblasts per brain lobe as compared to the 8 ± 0 type II neuroblasts consistently identified in wild-type brains (n = 10 brains per genotype; Figure 2A). Zld misexpression in all immature INPs driven by *Erm-Gal4* (*II*) resulted in 25.4 ± 8.7 type II neuroblasts per lobe (n = 10 brains). While fewer supernumerary type II neuroblasts (11.1 ± 2.0 type II neuroblasts per lobe; n = 10 brains) were identified when Zld misexpression was limited to late immature INPs and mature INPs (*Erm-Gal4* (*III*)), these data demonstrate that Zld misexpression can revert partially differentiated neuroblast progeny back to an undifferentiated stem-cell fate. Unlike Zld misexpression in immature INPs, expression of Zld in mature INPs driven by *Opa-Gal4* was not sufficient to induce supernumerary neuroblast formation (8.4 ± 0.5 type II neuroblasts per lobe; n = 8 brains). Thus, the ability of misexpression of Zld to promote the undifferentiated state is limited along the neuroblast lineage. This stands in contrast to aberrant Notch activation in mature INPs that can result in supernumerary neuroblasts. Thus, Zld promotes an undifferentiated state by functioning in parallel to, and not downstream of, Notch.

**Figure 2.**
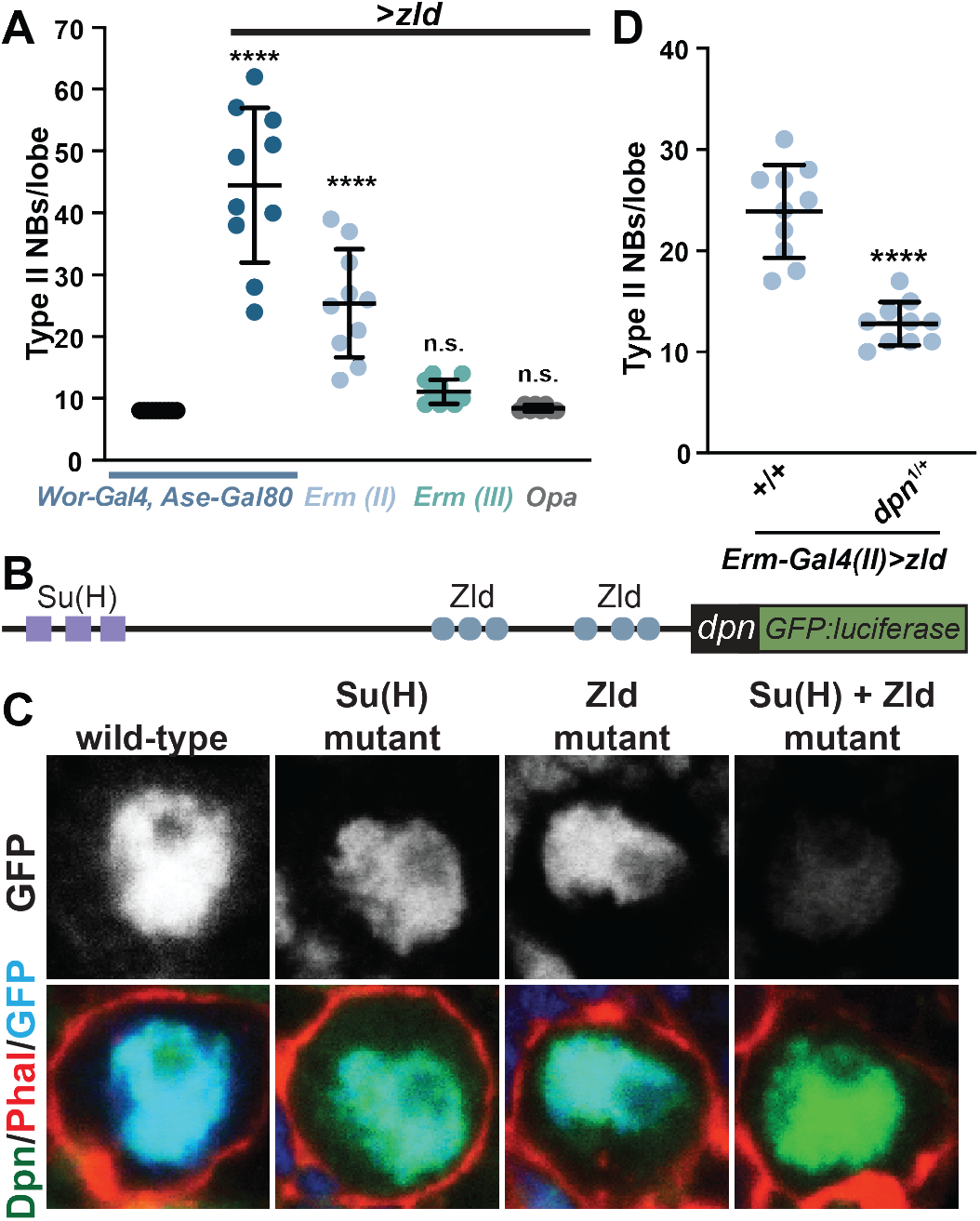
Zld promotes an undifferentiated state by activating *dpn* expression. A. Quantification of type II neuroblasts per lobe when Zld expression is driven by the indicated drivers (see Figure 1A for expression timing). Error bars show the standard deviation. *Wor-Gal4, Ase-Gal80* (8 ±0 n =10), *Wor-Gal4, Ase-Gal80 >zld* (44.5 ±12.5 n = 10, *Erm-Gal4(II) >zld* (25.4 ±8.7 n = 10, *Erm-Gal4*(*III*) >*zld* (11.1 ±2.0 n = 10, *Opa-Gal4* >*zld* (8.4 ± 0.5 n = 8). ‘ Significance is compared to *Wor-Gal4, Ase-Gal80* with a one-way ANOVA with post hoc Dunnett’s multiple comparisons test; n.s. = p-value > 0.05, **** = p-value ≤ 0.0001. B. Diagram of the *cis-*regulatory region of *dpn* used to create GFP:luciferase reporter constructs. Relative positions of Su(H)- and Zld-binding motifs are indicated. C. Representative images of GFP expression *(top)* in type II neuroblasts of animals expressing transgenes containing the wild-type binding sites, mutated Su(H)-binding sites, mutated Zld-binding sites, or mutated Su(H)- and Zld-binding sites. Staining for markers of the neuroblasts are shown *below*. D. Quantification of the number of type II neuroblasts per lobe upon exogenous expression of *zld* in immature INPs using the *Erm-Gal4(III)* driver in either wild type (23.9 ± 4.6) or *dpn^−/+^* (12.8 ± 2.1) larvae (n = 10). Comparison done using a two-tailed Student’s t test; **** = p-value ≤ 0.0001. See also Figure S2.

Notch functions by promoting expression of transcriptional repressors, including Dpn, to maintain type II neuroblasts in an undifferentiated state (Janssens et al., 2017; San-Juán and Baonza, 2011; Zacharioudaki et al., 2012; Zhu et al., 2012). However, Dpn remains expressed in type II neuroblasts even in the absence of Notch signaling, suggesting that additional activators can drive expression (San-Juán and Baonza, 2011). Zld binds to the *cis*-regulatory regions of *dpn* in the early embryo, and embryonic *dpn* expression depends on maternally encoded Zld (Figure S2A) (Harrison et al., 2011; Schulz et al., 2015). Thus, we hypothesized that Zld may promote an undifferentiated state in type II neuroblasts by functioning along with Notch to activate *dpn* expression.

To test the role of Zld in driving *dpn* expression, we generated transgenic reporters containing the *cis*-regulatory region of *dpn* driving GFP:luciferase (Figure 2B). This five kilobase region includes regulatory regions necessary for expression in both the embryo and neuroblasts, including several Zld-bound loci, as identified by ChIP-seq in the early embryo, and a cluster of previously identified binding sites for the Notch-binding partner Su(H) (Figure 2B) (Harrison et al., 2011; San-Juán and Baonza, 2011). In addition to the wild-type reporter, we created reporters with either the Zld-binding motifs, the Su(H)-binding motifs, or both Zld and Su(H)-binding motifs mutated. Similar to the endogenous locus, expression of the reporter depended on Zld binding for expression in the embryo (Harrison et al., 2011; Schulz et al., 2015) (Figure S2B). GFP expression was evident in type II neuroblasts of larva carrying the reporter with the wild-type *dpn*-regulatory region (Figure 2C). Mutation of either Su(H) or Zld-binding motifs reduced, but did not eliminate, GFP expression. Only mutation of both sets of binding motifs abrogated expression, demonstrating that *dpn* is a target of Zld in type II neuroblasts, similar to the embryo, and that Zld functions redundantly with Notch signaling to activate *dpn* expression.

Because of the ability of Zld and Notch to both activate *dpn* expression, to determine the functional significance of Zld-mediated activation of *dpn* expression we needed to test the effect of Zld expression in the absence of active Notch signaling. For this purpose, we focused on misexpression of Zld in immature INPs where Notch signaling is not active (Figure 1A). The supernumerary phenotype caused by Zld misexpression in immature INPs driven by *Erm-Gal4(II)* (23.9 ± 4.6 type II neuroblasts per lobe; n =10 brains) was strongly suppressed by loss of a single copy of *dpn* (12.8 ± 2.1 type ll neuroblasts per lobe; n = 10 brains; Figure 2D). These data demonstrate that Zld expression in immature INPs promotes reversion to an undifferentiated stem-cell fate at least in part by driving *dpn* expression. We note that while *Erm-Gal4(II)* is active in both immature INPs and mature INPs, this observed suppression is not due to reducing Zld-induced *dpn* expression in mature INPs, where Notch signaling is reactivated. This is because *dpn* overexpression in mature INPs cannot induce a supernumerary type II neuroblast phenotype. Thus, Zld functions in parallel to Notch to maintain type II neuroblasts in an undifferentiated state by activating *dpn* expression, and misexpression of Zld in the partially differentiated neuroblast progeny can revert them to an undifferentiated state by activating *dpn* expression.

### Zelda binds thousands of sites in type II neuroblasts

Having demonstrated that Zld promotes the undifferentiated type II neuroblast fate and that this activity is dependent, at least partially, on DNA binding, we used chromatin immunoprecipitation coupled with high-throughput sequencing (ChIP-seq) to identify Zld-binding sites in the type II neuroblasts. These experiments required enriching for type II neuroblasts, as wild-type larval brains only contain eight type II neuroblasts per lobe. For this purpose, we performed ChIP-seq on third instar larval brains dissected from larvae that are mutant for *brain tumor* (*brat^11/Df(2L)Exel8040^*). These *brat* mutant brains contain thousands of type II neuroblasts at the expense of other cell types and are a well characterized model for studying the transition from an undifferentiated stem-cell state to a committed INP identity (Janssens et al., 2017; Komori et al., 2018; Reichardt et al., 2018). Furthermore, these brains provide a biologically relevant system as the supernumerary neuroblasts are capable of differentiating along the type II lineage when the activity of genes that maintain neuroblasts in an undifferentiated state are inhibited (Rives-Quinto et al., 2020). Using the same Zld antibody previously used for Zld ChIP-seq in the early embryo (Harrison et al., 2011), ChIP-seq was performed in duplicate. The high correlation between replicates (Pearson’s correlation = 0.89) (Figure S3A), allowed us to identify 12,208 high-confidence peaks. Among the Zld-bound regions were the regulatory regions for genes known to maintain type II neuroblasts in an undifferentiated state, like *klumpfuss* (*klu*) (Berger et al., 2012; Xiao et al., 2012a) (Figure 3A). Identified Zld-binding sites were located in promoters and enhancers and were enriched for the known Zld-binding motif, CAGGTA (Figure 3B,C). Supporting the functional relevance of these Zld-binding sites, we identified robust peaks in the *dpn cis*-regulatory region that correspond to those regions mutated in our transgenic assays (Figure 4A). *De novo* motif enrichment also identified motifs known to be bound by additional proteins that have important functions in promoters as well as in three-dimensional chromatin organization (Van Bortle et al., 2014; Ohler et al., 2002; Ramírez et al., 2018; Zabidi et al., 2015) (Figure 3C).

**Figure 3.**
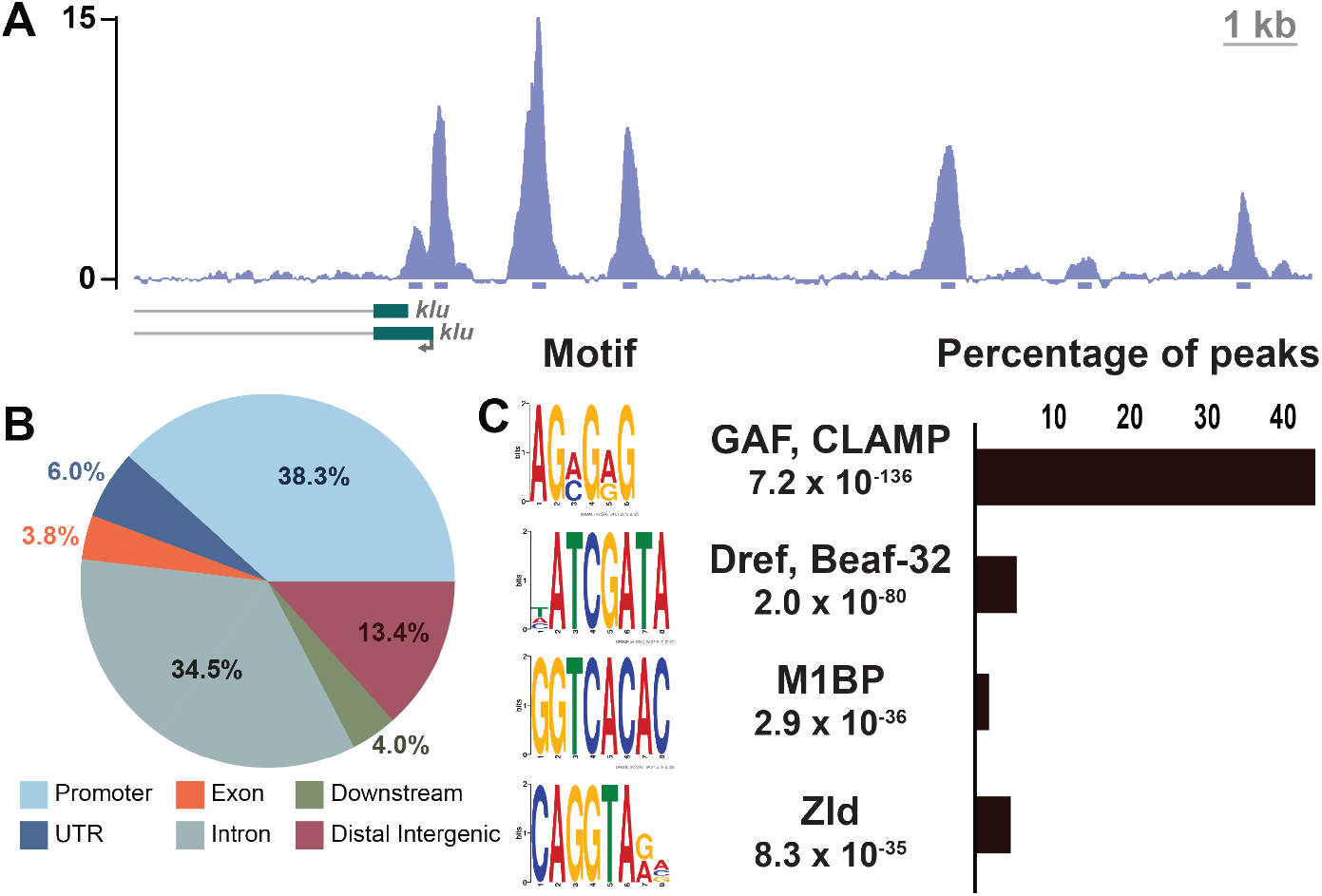
Zld binds to thousands of loci in type II neuroblasts. A. Representative z score-normalized genome browser tracks at the *klu* locus of Zld ChIP-seq from *brat^11/Df(2L)Excel8040^* brains. 200 bp regions surrounding the peak summits are shown below the track. B. Pie chart of the genomic distribution of Zld-binding sites in type II neuroblasts. Promoters are defined as −500 bp to +150 bp from the transcription start site. C. List of enriched motifs for Zld-bound regions in type II neuroblasts identified by MEME-suite (*left*). The significance value is based on the log likelihood ratio, width, sites, background, and size of the input sequence. Histogram of the fraction of the 12,208 peaks identified for Zld ChIP-seq containing the GAF, CLAMP (ACMGRG), Dref, Beaf-32 (HATCGATA), M1BP (GGTCACA) or Zld (CAGGTARV) motif (*right*). See also Figure S3.

### Zelda binds distinct sites in the type II neuroblasts and early embryo

In the early embryo, Zld binding is distinctive from that of other transcription factors. Zld binding is driven by DNA-sequence with 64% of the canonical Zld-binding motif (CAGGTAG) bound by Zld in the very early embryo (Harrison et al., 2011). Zld is a pioneer factor that can bind to nucleosomal DNA (McDaniel et al., 2019), and this capacity likely contributes to the unique binding profile in the early embryo. However, the naïve chromatin environment of the early embryo may also contribute to the unique binding profile of Zld at this stage. To determine the relative contributions of the pioneering activity of Zld and the naïve chromatin environment to the binding profile of Zld, we compared the binding of Zld in the type II neuroblasts to that of the early embryo. We realigned previously published ChIP-seq data for Zld from stage 5 embryos (Harrison et al., 2011) to the dm6 genome release, using the same parameters for aligning, filtering and peak calling as was used for the neuroblast ChIP-seq data. This allowed us to compare Zld-bound regions between the embryo and the larval type II neuroblasts. As might be predicted for a pioneer factor, we identified 4,269 regions that were bound by Zld at both stages of development, including the *dpn cis*-regulatory region (Figure 4A,B). Contrary to our expectations for a pioneer factor whose binding is driven strongly by sequence, we identified many more regions that were uniquely bound at a single stage of development (14,528 regions specifically bound in the stage 5 embryo and 7,939 regions specifically bound in the type II neuroblasts) (Figure 4B). These uniquely bound regions were not the result of differences in ChIP efficiency between the tissues as these regions include some of those with the highest relative peak height (Figure 4B). Because Zld binding in the embryo was strongly driven by sequence, the unique binding profile identified in the type II neuroblasts was unexpected. We therefore sought to confirm the identified Zld-binding sites using an additional antibody. For this purpose, we verified expression of our previously engineered endogenously sfGFP-tagged Zld line in the larval neuroblasts (Figure S3B) (Hamm *et al*., 2017) and used an anti-GFP antibody for ChIP-seq to identify binding of sfGFP-Zld in *brat* mutant brains. These experiments confirmed the cell type-specific Zld-binding profile (Figure S3C-F).

**Figure 4.**
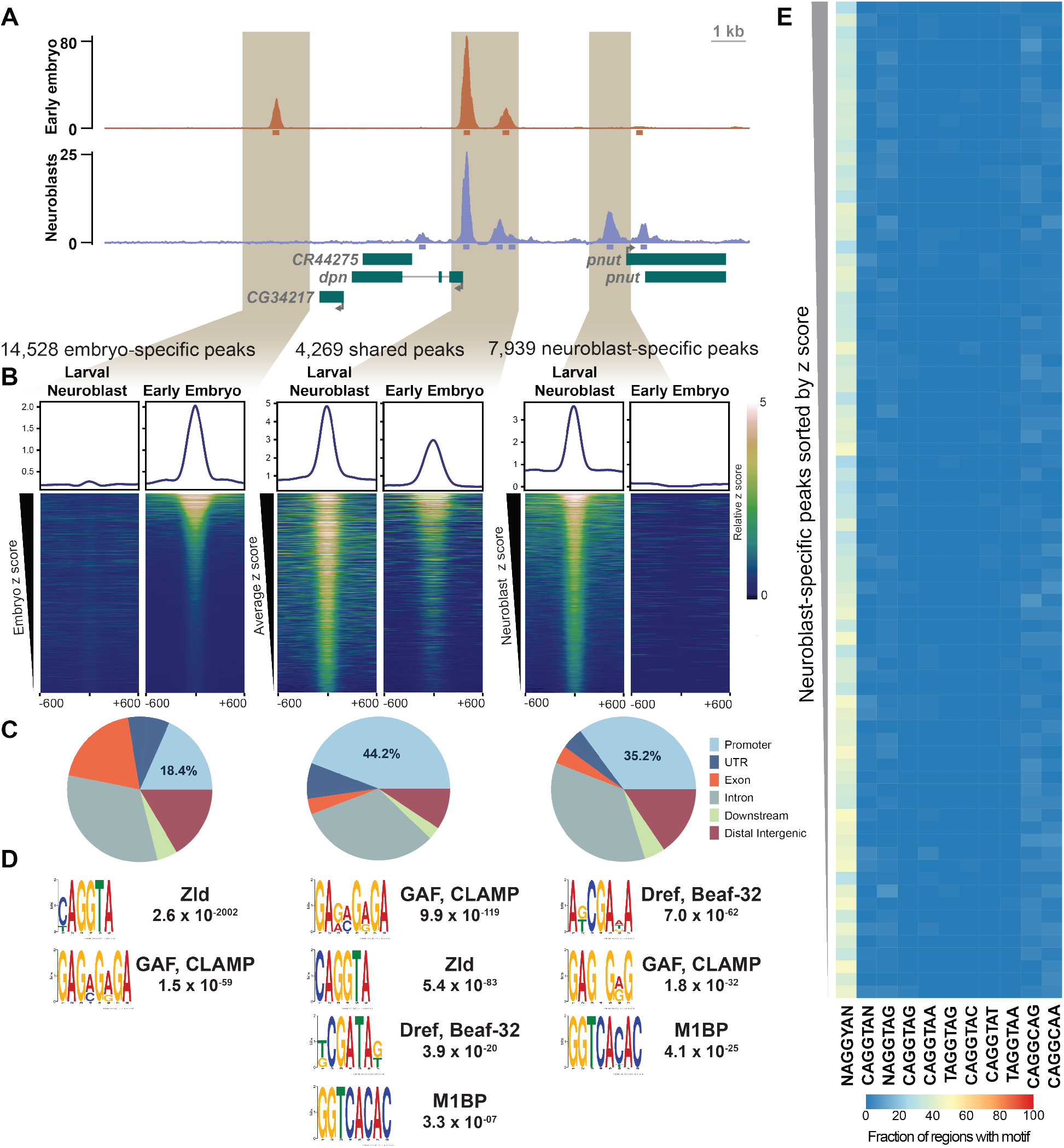
Zld occupancy in type II neuroblasts is driven by a feature apart from DNA sequence. A. Representative genome browser tracks for Zld ChIP-seq from stage 5 embryos (Harrison et al. 2011) and *brat^11/Df(2L)Excel8040^* brains centered on the *dpn* locus. 200 bp regions surrounding the peak summits are shown below the tracks. B. Heat maps centered on the ChIP peak summit with 600 bp flanking sequence for the embryo-specific peaks ranked by embryospecific peak signal, shared peaks ranked by the average signal across the embryo and type II neuroblasts, and type II neuroblast-specific peaks ranked by type II neuroblast-specific peak signal. Colors indicate relative ChIP-seq z score. Average z score profile is shown above each heat map. C. Pie charts of the genomic distribution of Zld-binding sites in each peak class. Promoters are defined as −500 bp to +150 bp from the transcription start site. D. Motif enrichment for each peak class as determined by MEME-suite. The significance value is based on the log likelihood ratio, width, sites, background, and size of the input sequence. E. Enrichment of Zld-binding motif variants among type II neuroblastspecific Zld peaks. Peaks are sorted based on z score from highest (top) to lowest (bottom) and binned into groups of 100. Color of each cell represents the percent of peaks within each bin containing the indicated motif. See also Figures S3-S4.

To determine what features shape Zld binding, we determined the genomic distribution of Zld-bound regions specific to the embryo, specific to the type II neuroblasts, and those shared between both cell types (Figure 4C). Regions bound by Zld in the type II neuroblasts, both those shared with the embryo and those unique to the neuroblasts, were enriched for promoters (44.2% and 35.2%, respectively; Figure 4C) as compared to regions bound by Zld solely in the embryo (18.4%; p-value < 2.2e-16) or randomized regions of the genome (10.4%; p-value < 2.2e-16) (Figure 4C, Figure S4A). Indeed, *de novo* motif analysis of Zld-bound regions identified multiple promoter-enriched sequence elements in these regions (Figure 4D). Similar to previous analysis of Zld-binding sites in the embryo, Zld-bound regions unique to the embryo were strongly enriched for the canonical Zld motif (Harrison et al., 2011) (Figure 4D). Unexpectedly, Zld-bound regions unique to the type II neuroblasts were not enriched for the Zld-binding motif (Figure 4D). Instead, motifs of known promoter-binding factors (Dref/Beaf-32, GAF/CLAMP and M1BP) were enriched (Figure 4D, Figure S4B). These factors have insulator function and bind in the promoters of housekeeping and constitutively active genes, where a large portion of chromatin boundaries are located (Van Bortle et al., 2014; Ohler et al., 2002; Ramírez et al., 2018; Zabidi et al., 2015). Indeed, when promoters were removed from the set of Zld-bound regions unique to the type II neuroblasts, *de novo* motif analysis did not enrich for these sequences, suggesting that the enrichment was due to Zld binding to promoters. Because bulk analysis can obscure sequences that might be enriched in a subset of bound regions, we binned Zld-bound regions by peak height and identified the fraction of regions in each bin that was enriched for the canonical Zld-binding motif or variants of this motif (Figure 4E, Figure S4C-F). As had been previously reported, Zld-bound regions in the embryo are highly enriched for the Zld-binding motif. By contrast, Zld-bound regions in the type II neuroblasts are only weakly enriched for a degenerate version of the canonical CAGGTA Zld-binding motif (Figure 4E). Together this analysis supports a model whereby in the embryo Zld binding is driven largely by sequence, but in the type II neuroblasts genomic features, apart from sequence, shape Zld binding.

### Chromatin accessibility shapes Zelda binding and function in type II neuroblasts

Because chromatin accessibility is known to influence the access of transcription factors to the underlying DNA, we assayed chromatin accessibility in brains enriched for type II neuroblasts using the Assay for Transposase-Accessible Chromatin using sequencing (ATAC-seq) (Buenrostro et al., 2016) (Figure S5A). We identified 58,551 accessible regions in the type II neuroblasts, including promoters of genes encoding factors that promote the maintenance of the undifferentiated state such as Dpn, Klu and Enhancer of split mγ (E(spl)mγ) (Figure S5B). 92% of loci bound by Zld in type II neuroblasts overlapped with regions of accessible chromatin as assayed by ATAC-seq, including both promoters and upstream *cis*-regulatory modules (Figure 5A). This correlation between Zld binding and open chromatin was similar to the previously reported association between Zld-bound regions and chromatin accessibility in the early embryo (Harrison et al., 2011). To better understand this relationship between accessibility and Zld binding, we used recently published ATAC-seq data generated in stage 5 embryos (Nevil et al., 2020) to allow us to compare accessibility and Zld binding in both the embryo and type II neuroblasts. This analysis showed that while Zld-bound loci and regions of open chromatin differ between the two cell types, there is a correlation between binding and accessibility in both (Figure 5A, Figure S5C). Regions bound by Zld only in the embryo are accessible in the embryo, but not the type II neuroblasts (Figure 5B). Similarly, regions bound specifically in the type II neuroblasts are more accessible in the neuroblasts than the embryo (Figure 5C).

**Figure 5.**
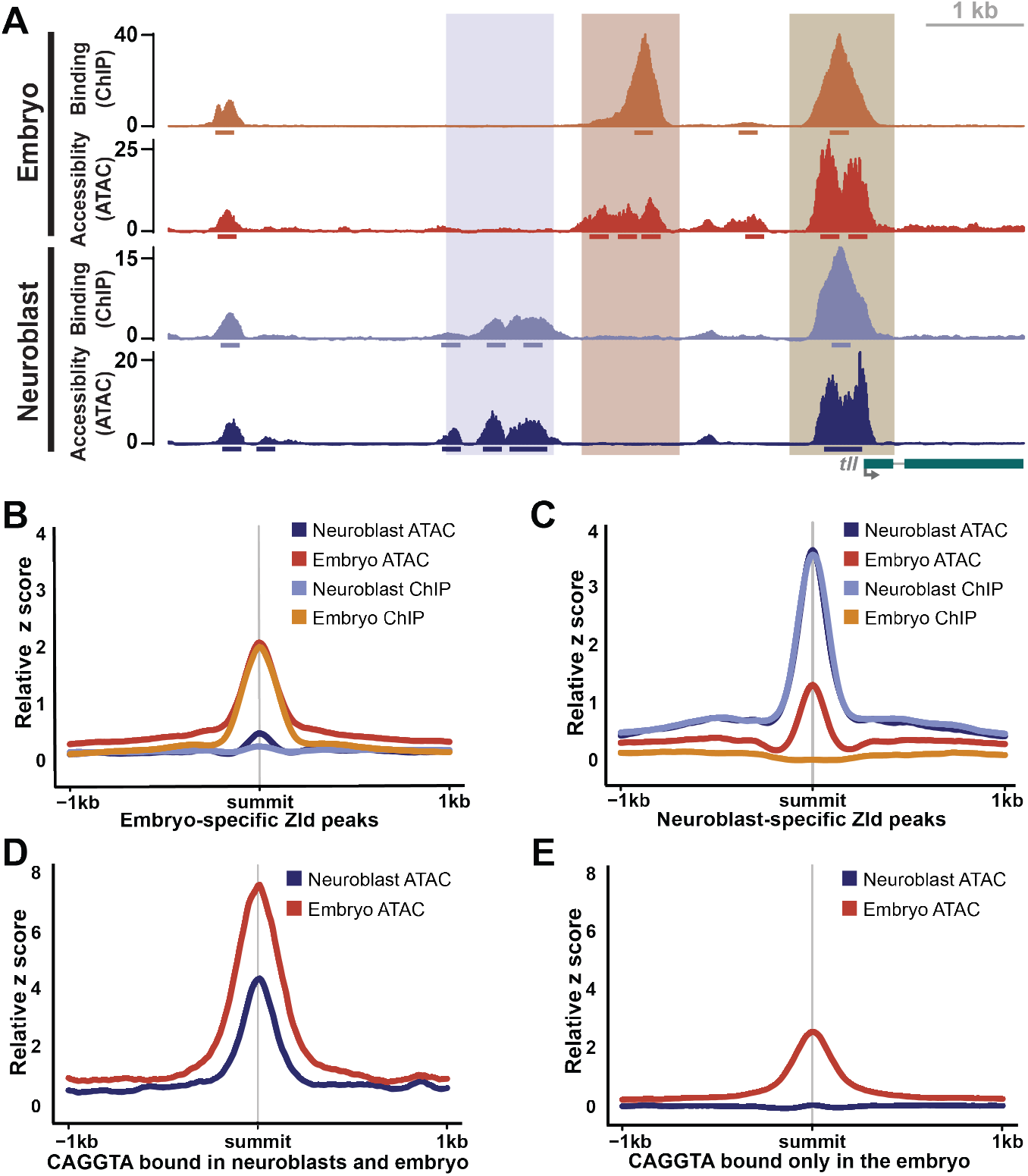
Cell type-specific differences in chromatin accessibility correlate with cell type-specific Zld binding. A. Genome browser tracks of Zld binding (ChIP-seq) and chromatin accessibility (ATAC-seq) in stage 5 embryos and type II neuroblasts of *brat* mutant brains at the *tll* locus. Peak regions are indicated below the tracks. Highlighted regions indicate regions bound by Zld and accessible only in neuroblasts (blue), only in embryos (orange) and in both (brown). B-C. z score for ChIP-seq and ATAC-seq centered on Zld ChIP-seq peaks specific to the embryo (B) or Zld ChIP-seq peaks specific to the type II neuroblasts (C). D-E. z score for ATAC-seq from the embryo and type II neuroblasts centered on the shared Zld ChIP-seq peaks containing a CAGGTA motif (D) or Zld ChIP-seq peaks specific to the embryo containing a CAGGTA motif (E). See also Figure S5.

At a subset of Zld-bound regions in the early embryo, Zld is required for chromatin accessibility (Schulz et al., 2015; Sun et al., 2015). Thus, the correlation between Zld binding and open chromatin might be due to Zld-mediated chromatin accessibility. However, chromatin accessibility also influences transcriptionfactor binding, such that accessibility may guide Zld binding to some loci. To distinguish between these two possibilities, we analyzed chromatin accessibility at regions containing the canonical Zld-binding CAGGTA motif. CAGGTA-containing regions bound in the type II neuroblasts and embryo are accessible in both cell types (Figure 5D). By contrast, CAGGTA-containing regions that are only bound in the embryo are only accessible in the embryo (Figure 5E). This demonstrates that in the type II neuroblasts Zld does not bind to regions containing the canonical Zld motif if the region is not accessible and suggests that the correlation between Zld binding and open chromatin in the neuroblasts may result from Zld occupying regions of accessible chromatin rather than Zld binding driving the accessibility.

### Zelda-binding sites define cell-type-specific enhancers

Our ChIP-seq analysis revealed thousands of Zld-binding sites that were unique to either the embryo or the type II neuroblasts (Figure 4A,B). By contrast, the majority of genes associated with Zld-bound regions in the type II neuroblasts were also associated with Zld binding in the embryo (Figure 6A, Table S1). This suggests that the transcriptional network regulated by Zld is partially shared between cell types. However, the identification of Zld-bound regions in neuroblasts lacking the canonical Zld-binding motif indicated that cell-type-specific constraints influence target recognition by Zld in non-embryonic cells and that the regulation of the shared gene network may depend on Zld binding to cell-type-specific genomic locations.

**Figure 6.**
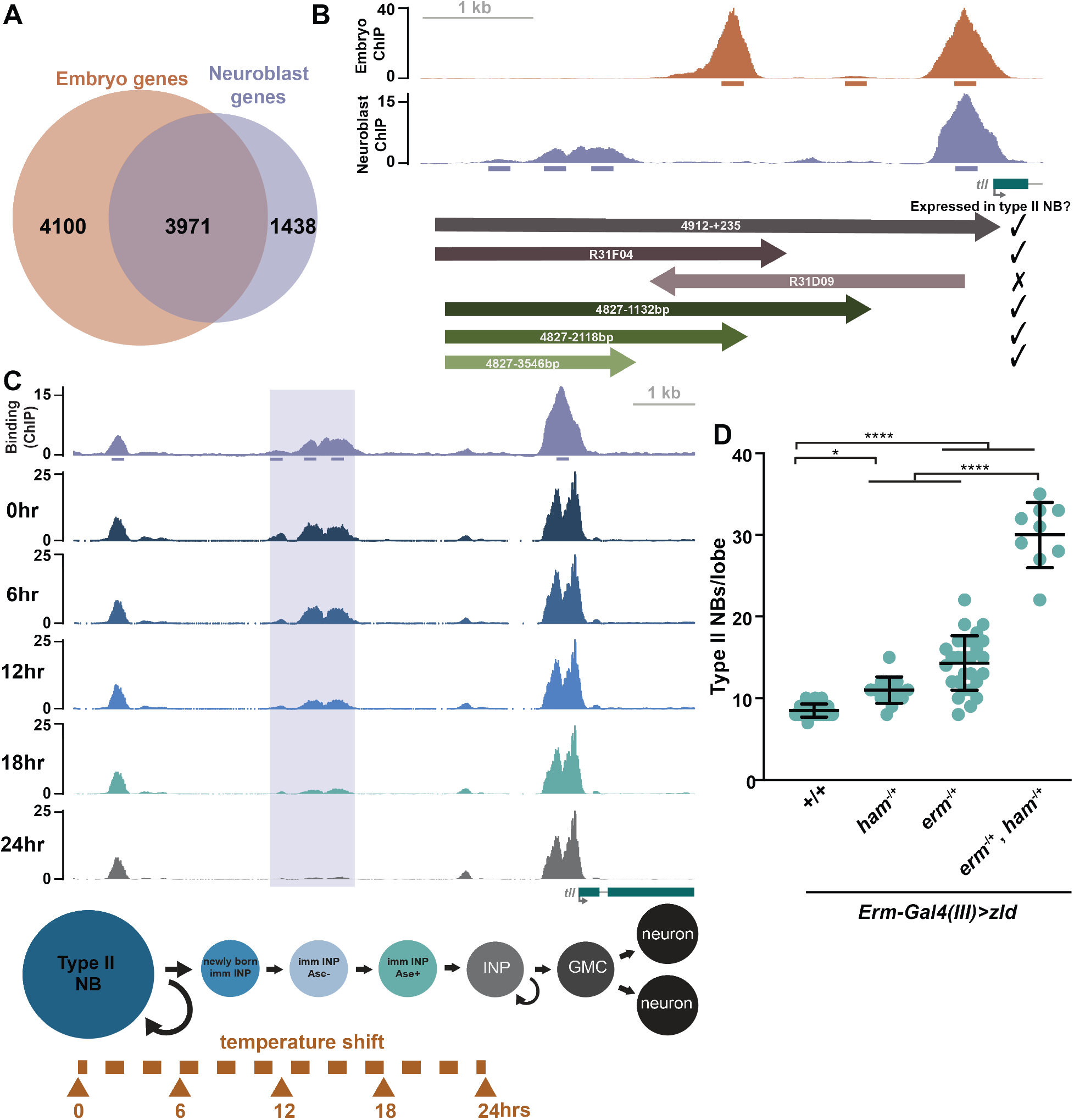
Erm and Ham limit Zld binding to a type II neuroblast-specific *tll* enhancer. A. Venn diagram of the genes associated with Zld ChIP-seq peaks from the early embryo and type II neuroblasts. B. Genome browser tracks of Zld binding (ChIP-seq) in stage 5 embryos and type II neuroblasts at the *tll* locus with arrows below indicating genomic regions tested for GFP expression in type II neuroblasts. Arrows show the relative direction of incorporation into transgenic reporters, and numbers indicate the sequence included relative to the transcription start site. C. Genome browser tracks of Zld ChIP-seq in type II neuroblasts of *brat* mutant brains and of ATAC-seq of the *tll* locus from *brat* mutant brains at the indicated time points following a temperature shift that initiates synchronous differentiation. Diagram below indicates approximates cell types enriched at each time point. 200 bp regions surrounding the ChIP peak summit are shown below the top track. D. Quantification of type II neuroblasts per lobe upon Zld expression in immature INPs *(Erm-Gal4(III))* in wild-type (8.5 ± 0.8) n = 23, *ham^−/+^* (11 ± 1.6) n = 15, *erm^−/+^* (14.3 ± 3.3 n = 28, *erm^−/+^, ham^−/+^* (30 ± 4.0) n = 9. Error bars show standard deviation. Comparisons were performed with a one-way ANOVA with post hoc Tukey’s HSD test; * = p-value ≤ 0.05, **** = p-value ≤ 0.0001. See also Figures S6-S7.

To begin characterizing changes that drive Zld binding to non-canonical sites, we focused on the gene *tll*, which is expressed in both the type II neuroblasts and the embryo (Hakes and Brand, 2020; Liaw and Lengyel, 1993; Rives-Quinto et al., 2020). Zld was bound to the *tll* promoter in both the embryo and type II neuroblasts, but Zld occupied different upstream regions at each developmental time point (Figure 5A). To determine if these upstream regions drive cell-type-specific gene expression, we analyzed reporter expression controlled by 5 kb of *tll* upstream-regulatory sequence, containing both the neuroblast-specific and embryo-specific Zld-bound regions. In embryos, this sequence can drive expression identical to the endogenous *tll* locus, and we showed it is sufficient to drive reporter expression in type II neuroblasts mimicking endogenous Tll expression (Figure 6B) (Liaw and Lengyel, 1993). Prior studies demonstrated that in the early embryo a transgene containing only 3 kb of upstream-regulatory sequence (and therefore lacking the neuroblast-specific Zld-binding sites), drives expression in a pattern identical to wild-type, but with reduced levels in the posterior (Liaw and Lengyel, 1993). Thus, the region bound by Zld specifically in the type II neuroblast is not necessary for expression in the early embryo, suggesting it may define a neuroblast-specific *cis*-regulatory module.

We more specifically tested if the type II non-canonical, neuroblast-specific Zld-binding sites were required to drive expression in the neuroblasts by examining expression of transgenes from the FlyLight collection containing portions of the *tll* regulatory region. R31F04, which contains sequence corresponding to the region extending from −4.9 to −1.8 kb upstream of the *tll* transcriptional start site, drove expression in the type II neuroblasts (Figure 6B). By contrast, a portion of the regulatory region that includes the embryo-specific Zld-bound region, but not the type II neuroblast-specific region (FlyLight R31D09) failed to drive expression in the type II neuroblasts. This construct is inserted in the reverse orientation compared to the endogenous locus, and, while enhancers generally function regardless of genomic orientation, it remains possible that this difference in orientation is what results in the lack of expression. Together these data further suggest that the region bound by Zld specifically in neuroblasts constitutes an enhancer required for *tll* expression in type II neuroblasts.

We were unable to identify and mutate the sequence that was necessary for Zld binding, as was done for the *dpn* regulatory region, because the region underlying the type II neuroblast-specific Zld-binding sites does not contain the canonical Zld-binding motif. We therefore created three transgenes in which sequentially truncated portions of the *tll* upstream regulatory sequence drove expression from the *Drosophila* synthetic core promoter element (DSCP) and assayed for expression in the type II neuroblasts (Figure 6B). All three fragments tested were sufficient to drive reporter expression in type II neuroblasts (Figure 6B, Figure S6). Together these assays show that type II neuroblast- and embryo-specific binding by Zld identifies enhancers required for *tll* expression specifically in each cell type.

In addition to having cell-type-specific Zld occupancy, these upstream regions also show celltype-specific patterns of chromatin accessibility (Figure 5A) as might be predicted for cell-type-specific enhancers. In the type II neuroblast lineage, *tll* is expressed exclusively in the neuroblast and not in other cell types. Indeed, misexpression of *tll* in mature INPs results in their reversion to neuroblasts (Hakes and Brand, 2020; Rives-Quinto et al., 2020). To investigate whether chromatin accessibility at the Zld-bound region reflects this expression pattern, we took advantage of a synchronous, time-release differentiation system we developed that recapitulates many of the gene expression changes that occur as type II neuroblasts differentiate into INPs (Rives-Quinto et al., 2020) and performed ATAC-seq at four time points spanning the first 24 hours following temperature shift (approximating differentiation from type II neuroblast to INP). Consistent with Tll expression rapidly diminishing in immature INPs, the region with the most dramatic loss in accessibility over this time course was the Zld-bound, type II neuroblast-specific *tll* enhancer (Figure 6C).

Like *tll, Six4* expression significantly decreased in our synchronous, time-release differentiation system (Rives-Quinto et al., 2020). *Six4* encodes a homeodomain-containing transcription factor with known roles mesoderm specification (Clark et al., 2006; Kirby et al., 2001). Using GFP reporter expression, we confirmed that *Six4* is expressed in type II neuroblasts and not in INPs (Figure S7A) (Chen et al. 2021). We further identified a neuroblast-specific Zld-bound region upstream of *Six4* that significantly lost chromatin accessibility upon neuroblast differentiation in our time-release system (Figure S7B). Together these data support a model in which binding of the pioneer factor Zld in neuroblasts defines enhancers that drive type II neuroblast-specific expression of *tll* and *Six4*, and theses enhancers progressively lose accessibility as expression is silenced and cells differentiate.

We have recently shown that Erm and Ham function through Hdac3 to silence *tll* expression during INP commitment, thus preventing aberrant *tll* expression in mature INPs following reactivation of Notch signaling (Rives-Quinto et al., 2020). The timing of Erm- and Ham-mediated silencing of *tll* coincides with the loss of chromatin accessibility at the Zld-bound, type II neuroblast enhancer (Figure 1A). Thus, it is possible that the chromatin changes implemented by Erm and Ham limit the ability of Zld to bind this enhancer. If misexpression of Zld in INPs is sufficient to drive Tll expression, then Zld misexpression would be expected to mimic Tll misexpression in these cells and promote supernumerary type II neuroblast formation (Hakes and Brand, 2020; Rives-Quinto et al., 2020). However, Zld expression in INPs (driven by *Opa-Gal4*) failed to induce supernumerary type II neuroblast formation, suggesting cell-type-specific features limit the ability of Zld to activate *tll* expression (Figure 2A). To determine if the sequential silencing of *tll* by Erm and Ham limits the ability of Zld to induce INP reversion to type II neuroblasts, we drove Zld expression in immature INPs and mature INPs (*Erm-Gal4 (III)*) of brains heterozygous for null mutations in either *erm* or *ham*. While loss of single copies of either *erm* or *ham* does not result in supernumerary type II neuroblasts (Rives-Quinto et al., 2020), loss of a single copy of either of these genes enhanced the weak supernumerary phenotype caused by misexpression of Zld (14.3 ± 3.3 type II neuroblasts; n = 28 and 11 ± 1.6 type II neuroblasts; n = 15 respectively). This effect was further enhanced when copies of both *erm* and *ham* were removed (30 ± 4.0 type II neuroblasts; n = 9) (Figure 6D). Together these data support a model in which type II neuroblast-specific binding of the pioneer factor Zld defines an enhancer that drives expression of a master regulator of type II neuroblast functional identity, Tll, and that this enhancer is progressively silenced by Erm and Ham to inhibit reactivation in the INP. Furthermore, we propose that other neuroblast-specific Zld-binding sites, such as those upstream of *Six4*, may define additional enhancers driving stem-cell fate in type II neuroblasts.

## Discussion

Our results demonstrate that Zld, an essential transcriptional activator of the zygotic genome, promotes the undifferentiated state in the neural stem cell lineage of the larva and can revert partially differentiated cells to a stem cell. Other pioneer factors are known to have similar functions when mis-expressed, and this can lead to disease. For example, expression of DUX4, an activator of the zygotic genome in humans, in muscle cells leads to Facioscapulohumeral muscular dystrophy (FSHD), and OCT4 and Nanog are overexpressed in undifferentiated tumors and their expression is associated with poor clinical outcomes (Ben-Porath et al., 2008; Chew et al., 2019; Hendrickson et al., 2017; Lemmers et al., 2010; Meng et al., 2010; Nagata et al., 2014; Whiddon et al., 2017). Despite the ability of Zld to promote the undifferentiated stem-cell fate, this capacity is limited as cells differentiate to INPs. We showed that the ability of Zld to drive gene expression is limited by changes to chromatin that are mediated by the repressors Erm and Ham. Because of the ability of pioneer factor misexpression to cause disease, understanding these cell type-specific constraints on pioneer factors has important implications for our understanding of development and disease.

### Zld functions in parallel to Notch to define type II neuroblast enhancers and promote an undifferentiated state

Zld expression promotes the reversion of partially differentiated immature INPs to a stem-cell fate resulting in supernumerary type II neuroblasts. Furthermore, failure to down-regulate *zld* in the newly generated INPs results in supernumerary type II neuroblasts (Reichardt et al., 2018). Thus, Zld levels must be precisely controlled to allow differentiation following asymmetric division of the type II neuroblasts. Zld promotes the undifferentiated state, at least in part, through the ability to drive expression of Dpn, a key transcription factor for driving type II neuroblast self-renewal (Bier et al., 1992; San-Juán and Baonza, 2011; Zacharioudaki et al., 2012, 2016; Zhu et al., 2012). *dpn* is a target of the Notch pathway in type II neuroblasts and constitutively activated Notch signaling drives *dpn* expression (San-Juán and Baonza, 2011; Zacharioudaki et al., 2016). However, loss of Notch signaling does not completely abrogate expression of known target genes, including *dpn*, suggesting that additional activators can drive expression in the absence of Notch (San-Juán and Baonza, 2011; Zacharioudaki et al., 2016; Zhu et al., 2012). Indeed, we show that Zld functions as such a factor in driving *dpn* expression and loss of a single copy of *dpn* can suppress the ability of Zld to promote the reversion of immature INPs to type II neuroblasts. We propose that this redundancy with Notch is not limited to regulating *dpn* expression and that Zld and Notch may function together to regulate a number of genes required for type II neuroblast maintenance.

Supporting this, Zld is bound to 49% of the identified direct Notch-target genes in neuroblasts (Table S1) (Zacharioudaki *et al*., 2016). Although, Zld is not required for type II neuroblast maintenance, loss of Zld can enhance knockdown of the Notch pathway demonstrating a partially redundant requirement for these two pathways in maintaining type II neuroblast fate. Based on these data, we propose that Zld and Notch function in parallel to drive gene expression, and this redundancy robustly maintains the type II neuroblast pool (Figure 7A).

**Figure 7.**
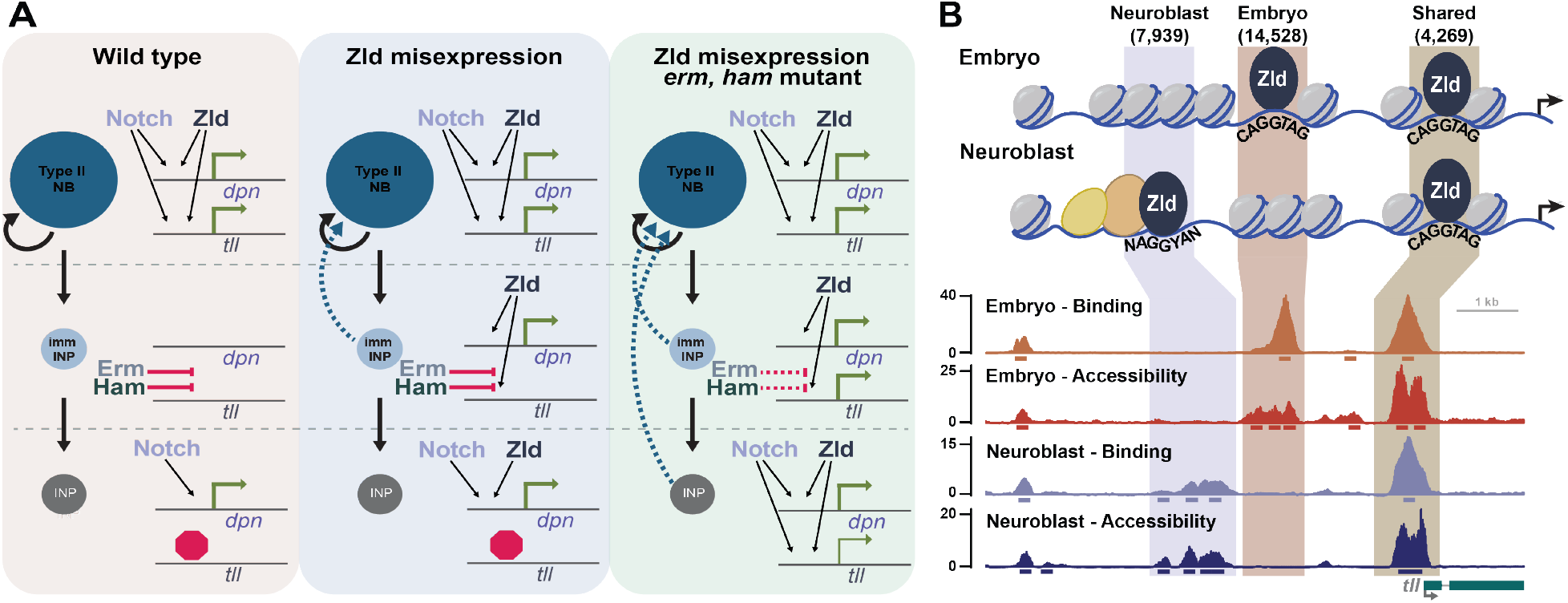
Zld promotes the undifferentiated state by defining neuroblast-specific enhancers that become progressively silenced during differentiation. A. Zld functions in parallel to Notch to maintain type II neuroblasts in an undifferentiated state through activation of *tll* and *dpn*. Downregulation of Notch signaling in newly born INPs allows Erm and Ham to become sequentially activated during INP commitment. In the INPs, Erm- and Ham-mediated silencing of *tll* (indicated by the red octagon/stop sign) prevents reactivation of Notch signaling from inducing *tll* expression and driving reversion to neuroblasts. When misexpressed in immature INPs, Zld activates *dpn* expression and promotes reversion to a neuroblast. Changes to the chromatin structure mediated by Erm and Ham limit the ability of Zld to drive *tll* expression and therefore reprogram mature INPs. B. Cell-type specific binding by Zld correlates with chromatin accessibility and defines tissue-specific enhancers. Type II neuroblast-specific Zld-binding sites are not enriched for the canonical Zld-binding motif and are instead enriched for sequences corresponding to additional cofactors that may stabilize Zld binding or promote chromatin accessibility at these loci.

### Chromatin occupancy by Zld is cell type-specific

Zld binding in the early embryo is distinctive as it is driven primarily by DNA sequence with a majority of the canonical-binding motifs occupied. This is in contrast to most other transcription factors, whose binding is influenced widely by chromatin accessibility and therefore bind only small fraction of their canonical motifs (Clark and Felsenfeld, 1971; Luger et al., 2012; Tremethick, 2007). Here we report, for the first time, the genome-wide occupancy of Zld in a tissue apart from the early embryo and begin to identify important functions for zygotically expressed Zld in a stemcell population. While we identified thousands of loci that are occupied by Zld both in the embryo and in the larval type II neuroblasts, thousands more were unique to each cell type. This is in contrast to what we have shown for another pioneer-transcription factor, Grainy head (Grh), which has similar genomic occupancy in the embryo and larval imaginal discs (Nevil et al., 2017, 2020). Thus, unlike for Grh, Zld binding is cell type-specific and likely governed by changes to the chromatin structure along with the expression of cell type-specific transcription factors (Figure 7B).

Despite their ability to engage nucleosomal DNA, experiments, largely in cell culture, have demonstrated that most pioneer transcription factors show cell-type specific chromatin occupancy (Buecker et al., 2014; Chronis et al., 2017; Donaghey et al., 2018; Mayran et al., 2018; Swinstead et al., 2016). Both chromatin state and co-factors influence the binding of pioneer factors, like Oct4 (Buecker et al., 2014; Chronis et al., 2017; Soufi et al., 2012, 2015). While most pioneer factors have been studied through misexpression in culture, our data show that binding of an endogenously expressed pioneer factor within a developing organism is also cell-type specific. By analyzing genome occupancy and chromatin accessibility in two different cell types, the embryo and type II neuroblast, we demonstrate that binding is highly correlated with accessibility in both cell types. While it is not possible to determine whether chromatin accessibility regulates Zld binding or Zld binding drives accessibility, we propose that in the larval type II neuroblasts accessibility influences Zld occupancy. In the early embryo, Zld binds when the chromatin is naïve with relatively few chromatin marks and is rapidly replicated (Hamm and Harrison, 2018). This Zld binding is driven largely by DNA sequence and is required for chromatin accessibility at a subset of sites (Harrison et al., 2011; Nien et al., 2011; Schulz et al., 2015). Thus, in the early embryo, Zld binding can influence accessibility. However, Zld occupancy is reorganized in the type II neuroblasts such that only a fraction of the canonical binding motifs is occupied, and those motifs that are not bound by Zld are not accessible. Furthermore, the type II neuroblast-specific binding sites are enriched at promoters, which are known to be generally accessible in a broad range of cell types. This suggests that Zld occupancy in the type II neuroblasts is likely shaped by chromatin accessibility. Together, these data support a model whereby in the early embryo Zld can bind broadly to the naïve genome while in the neuroblasts Zld binding is limited by chromatin state. Future studies will enable the identification of what limits Zld binding and will allow for the definition of chromatin barriers to reprogramming within the context of a developing organism.

Along with chromatin structure, cofactors regulate mammalian pioneer-factor binding in culture, including binding by Oct4, Sox-2 and FOXA2 (Buecker et al., 2014; Donaghey et al., 2018; Liu and Kraus, 2017). In addition to chromatin accessibility, our data similarly support a role for specific transcription factors in regulating Zld binding in type II neuroblasts. Type II neuroblast-specific Zld-bound loci are not enriched for the canonical Zld motif, suggesting additional factors facilitate Zld binding to these regions. Supporting a functional role for this recruitment, expression of Zld with mutations in the zinc-finger DNA-binding domain, which abrogate sequence-specific binding (Hamm et al., 2015, 2017), can still drive supernumerary neuroblasts. We have previously shown that while this mutant protein lacks sequence-specific binding properties, the polypeptide retains an affinity for nucleosomal DNA (Fernandez Garcia et al., 2019; McDaniel et al., 2019). This non-specific affinity may be stabilized by additional factors expressed in the neuroblasts.

### Changes to the chromatin state limit Zld-mediated activation of neuroblast-specific enhancers

The capacity of Zld to drive the undifferentiated state is limited along the type II neuroblast lineage. While expression of Zld in immature INPs results in supernumerary neuroblasts, Zld expression in mature INPs does not. Similarly, misexpression of the Zld-target gene *dpn* in immature INPs can drive their reversion to neuroblasts, and our data suggest that Zld-mediated reversion is caused, at least in part, by driving expression of *dpn*. By contrast, the endogenous expression of *dpn* in mature INPs does not cause their reversion to neuroblasts because the self-renewal program is decommissioned during INP maturation. This decommissioning is mediated by successive transcriptional repressor activity (Rives-Quinto et al., 2020). We recently demonstrated that Erm and Ham function sequentially to repress expression of genes that promote neuroblast fate. Our data suggest that changes to the chromatin structure mediated by these transcriptional repressors limit the ability of Zld to drive gene expression and therefore reprogram mature INPs (Figure 7A).

An essential target of Erm and Ham repression is *tll* (Rives-Quinto et al., 2020). In contrast to other stem-cell regulators like Notch and Dpn, Tll is expressed only in type II neuroblasts and not in the transit-amplifying INPs (Hakes and Brand, 2020; Rives-Quinto et al., 2020). Furthermore, expression of *tll* in mature INPs can robustly drive supernumerary neuroblasts (Hakes and Brand, 2020; Rives-Quinto et al., 2020). In the embryo, *tll* is a Zld-target gene (McDaniel et al., 2019; Nien et al., 2011), and our ChIP-seq data identify Zld-binding sites in the type II neuroblasts. While Zld occupies the promoter of *tll* in both type II neuroblasts and the embryo, we identify unique binding sites for Zld in upstream regions in both cell types and demonstrate that these likely define cell type-specific enhancers. Our data identify a neuroblast-specific, Zld-bound enhancer that drives expression specifically in the neuroblasts and supports a model whereby Zld activates expression from this enhancer in the type II neuroblasts. Erm- and Ham-mediate chromatin changes, likely through histone deacetylation, that progressively limit chromatin accessibility during INP maturation. This decrease in accessibility inhibits the ability of ectopically expressed Zld to activate expression from this enhancer, keeping *tll* repressed in the INPs. Gene expression profiling identified *Six4* as a gene that, like *tll*, is expressed only in neuroblasts and not in INPs (Rives-Quinto et al., 2020). Here we identified a neuroblast-specific Zld-bound region that progressively loses accessibility during INP maturation. Thus, Erm and Ham likely silence multiple Zld-bound enhancers to allow for the transition from a self-renewing neuroblast to a transient-amplifying INP. We propose that in type II neuroblasts Zld is instructive in defining chromatin accessibility for this enhancer and that following asymmetric division, changes to the chromatin state mediated by Erm and Ham and down-regulation of Zld expression robustly decommissions this enhancer, allowing for differentiation (Figure 7).

Our data identify Erm- and Ham-mediated changes to the chromatin that inhibit binding by the pioneer factor Zld and, in so doing, limit the ability of Zld expression to reprogram cell fate. Our studies in both the early embryo and in type II neuroblasts provides a powerful platform for identifying the barriers to pioneer factor-mediated reprogramming within the context of development and support a role for both chromatin organization and cell-type specific co-factors in determining Zld occupancy. Future studies will reveal these specific barriers and in doing so will help to identify fundamental processes that may limit reprogramming both in culture and in disease states.

## Materials and Methods

### Drosophila strains

Flies were raised on standard fly food at 25°C (unless otherwise noted). To obtain the *brat* mutant brains used for ChIP-seq and ATAC-seq, we crossed *brat^11^/CyO, ActGFP* to *brat^Df(2L)Exel8040^/CyO, ActGFP* (BDSC#7847) (Arama et al., 2000; Lee et al., 2006a) and screened for GFP negative larvae. Imaging experiments took advantage of a line with endogenous Zld tagged with superfolder GFP (Hamm et al., 2017). For ChIP-seq experiments using the anti-GFP antibody, the sfGFP-tagged Zld alleles were combined with *brat^11^* and *brat^Df(2L)Exel8040^* and crossed to obtain *brat* mutant brains. Imaging of Six4::GFP (BDSC#67733) and FlyLight GMR31F04-Gal4 (BDSC#46187) and GMR31D09-Gal4 (BDSC#49676) was done with previously published lines.

For ectopic Zld expression experiments, the open reading frame for Zld-RB was cloned into the pUASt-attB vector using standard PCR, restriction digest and ligation procedures. Similarly, transgenic UAS driven Zld ZnF mutants were cloned using previously generated plasmids (Hamm et al., 2015) and a gene block containing ZnF mutations in the DBD of Zld (Integrated DNA Technologies, Coralville, IA). The mutated C2H2 ZnFs convert cysteines to serines. These transgenes were integrated at ZH-86Fb site on chromosome 3 using φC31-mediated integration (Bischof and Basler, 2008) (BestGene, Chino Hills, CA). *Worniu-Gal4; Asense-Gal80* (Neumuller et al., 2011), *Worniu-Gal4; Tub-Gal80ts* (Lee et al., 2006a), *Erm(II)-Gal4* (Xiao et al., 2012), *Erm(III-Gal4)* (Pfeiffer et al., 2008) and *Opa-Gal4* (BDSC#46979, *GMR77B05-Gal4*) drivers were used to drive ectopic expression.

To create the *dpn* transgenes we ordered gene blocks (Integrated DNA Technologies, Coralville, IA) with mutated Zld and Su(H) binding sites in the 5kb *dpn cis*-regulatory and used Gateway LR Clonase (ThermoFisher Scientific, Waltham, MA) to clone the region into the VanGlow vector without the DSCP (Addgene#83342) (Janssens et al., 2017). The various transgenic reporters were integrated into the VK37 site on chromosome 2 using φC31-mediated integration (Bischof and Basler, 2008) (BestGene, Chino Hills, CA). L3 brains from reporter fly lines were dissected, fixed and stained to visualize GFP reporter expression in the type II NBs. Embryos from these lines were collected on molasses plates with yeast paste from flies reared in cages for luciferase assays. The embryos were ground in an eppendorf tube with a pestle in 1x passive lysis buffer and luciferase signal was measured using the Dual Luciferase assay from Promega (Promega, Madison, WI). *tll* transgenes were similarly cloned with Gateway LR Clonase (ThermoFisher Scientific, Waltham, MA) to insert PCR-amplified genomic regions into the VanGlow vector with the DSCP (Addgene#83338). The various transgenic reporters were integrated into the VK31 site on chromosome 3 using φC31-mediated integration (Bischof and Basler, 2008) (BestGene, Chino Hills, CA).

*dpn^1^* (Bier et al., 1992), *ham^Df(2L)Exel7071^* (BDSC#7843) (Rives-Quinto et al., 2020) and *erm^l(2)5138^* (Weng et al., 2010) mutant experiments were done using previously published lines. *zld^294^* (Liang et al., 2008) and Notch RNAi (BDSC#33611) clones were generated using previously published methods (Lee et al., 2000). Briefly, clones were induced by heat shock at 37°C for 90 minutes at 24 hours after larval hatching. Brains were dissected for clone analysis at 72 hours after clone induction.

### Immunofluorescent staining and antibodies

Larval brains were dissected in PBS and fixed in 100 mM Pipes (pH 6.9), 1 mM EGTA, 0.3% Triton X-100 and 1 mM MgSO4 containing 4% formaldehyde for 23 minutes. Fixed brain samples were washed with PBST containing PBS and 0.3% Triton X-100. After removing fix solution, samples were incubated with primary antibodies for 3 hours at room temperature. At 3 hours later, samples were washed with PBST and then incubated with secondary antibodies for overnight at 4°C. The next day samples were washed with PBST and then equilibrated in ProLong Gold antifade mountant (ThermoFisher Scientific, Waltham, MA). The confocal images were acquired on a Leica SP5 scanning confocal microscope (Leica Microsystems Inc, Buffalo Grove, IL). More than 10 brains per genotype were used to obtain data in each experiment. Rabbit anti-Ase Antibody (1:400 for IF) (Weng et al., 2010). Rat anti-Dpn Antibody (clone 11 D1BC7.14; 1:2 for IF) (Lee et al., 2006b). Chicken anti-GFP Antibody (Aves Labs, Davis, CA, Cat #GFP-1020). Rhodamin phalloidin (ThermoFisher Scientific, Waltham, MA, Cat #R415).

### Chromatin Immunoprecipitation

500 brains (in 45 min time windows) were dissected in Schneider’s medium (Fisher, Hampton, NH, Cat #21720001) from *brat^11/Df(2L)Exel8040^* larvae aged for 5-6 days at 25°C (L3 stage). The dissected brains were fixed in 1.8% formaldehyde for 20 min, which was stopped by incubation with 0.25M Glycine at room temperature for 4 min and on ice for 10 min. The samples were washed 3 times with wash buffer (1xPBS, 5mM Tris-HCl pH 7.5, 1mM EDTA), flash frozen in liquid nitrogen and stored at −80°C until all the brains had been collected. Brains were thawed on ice and combined for homogenization in SDS lysis buffer (1% SDS, 1mM PMSF, 50mM Tris-HCl pH 8.1,5mM EDTA) containing protease inhibitors (Pierce mini tablets EDTA-free, VWR, Radnor, PA, Cat #PIA32955) to obtain nuclear extracts. The nuclear extracts were disrupted using a Covaris sonicator (S220 High Performance Ultrasonicator) (18 cycles of 170 Peak Power, 10 Duty Factor, 200 cycles/burst for 60 sec). 7% of the sonicated sample was stored as input and the rest was diluted with 1X volume of dilution buffer (1% Triton X-100, 280mM NaCl) and incubated overnight with antibodies (8μl anti-Zld (Harrison et al., 2011) or 1.4μl anti-GFP (Abcam, Cambridge, UK, Cat #ab290)). Protein A beads (Dynabeads Protein A, ThermoFisher Scientific, Waltham, MA, Cat #10002D) were added and samples were incubated at 4°C for 4 hrs. Beads were recovered and washed. (1x with low salt wash buffer (0.1% SDS, 1% Triton X-100, 2mM EDTA, 20mM Tris-HCl pH 8.1 and 150 mM NaCl), 1x high salt wash buffer (0.1% SDS, 1% Triton X-100, 2mM EDTA, 20mM Tris-HCl pH 8.1, 500mM NaCl), 1x LiCl wash buffer (0.25M LiCL, 1% NP40, 1% deoxycholate, 1mM EDTA, 10mM Tris-HCl pH8.1) and 2x with TE buffer (10mM Tris-HCl pH 8.0, 1mM EDTA).) Washed beads were incubated at room temperature for 15 min with elution buffer (0.1M NaHCO3, 1% SDS) to elute the chromatin. The samples and corresponding input were incubated at 55°C overnight with PK solution (15μl 1M Tris pH 7.5, 7μl 0.5M EDTA, 4μl 5M NaCl, 2μl 20mg/ml Proteinase K (Fisher, Hampton, NH, Cat #EO0491)). Samples were incubated with 0.5μl 20mg/ml RNase (PureLink RNase A, Thermo Fisher, Waltham, MA, Cat #12091021) at 37°C for 30 min and moved to 65°C for 6 hrs to reverse the crosslinking. Samples were cleaned up by phenol:chloroform extraction, precipitated, and resuspended in 20μl TE buffer. Sequencing libraries were made using the NEB Next Ultra II DNA library kit (New England BioLabs Inc, Ipswich, MA, Cat #E7645S) and were sequenced on the Illumina Hi-Seq4000 using 50bp single-end reads at the Northwestern Sequencing Core (NUCore).

### ChIP-seq Data Analysis

Read quality was checked using FASTQC (Andrews, 2010). Adapters and low-quality bases were removed using Trimmomatic-0.39. (Bolger et al., 2014). Reads were mapped to the dm6 genome assembly (Dos Santos et al., 2015) using Bowtie 2 (Langmead and Salzberg, 2012). Unmapped, multiply aligning, mitochondrial, and scaffold reads were removed. Throughout SAMtools was used to filter and convert file formats (Li et al., 2009). MACS version 2 (Zhang et al., 2008) was used with default parameters to identify bound regions of chromatin in samples (IP vs INPUT) for both replicates of Zld antibody ChIP in neuroblasts. The Zld antibody ChIP in the embryo did not have a corresponding INPUT for the single IP, so peaks were called without reference to a control. The GFP antibody ChIP in the neuroblasts was called with the parameters above with the exception of lowering the m-fold value to −m 3 50 due to low IP efficiency. Peak summits were extended by 100bps on either side. High confidence Zld-bound regions in the neuroblasts were called as 200bp peak regions with 50% overlap in both replicates using the BEDtools intersect function (Quinlan and Hall, 2010). Regions belonging to contigs and unmapped chromosomes were removed. High-confidence regions used for analysis with the GFP antibody ChIP in neuroblasts were called as being bound with 50% overlap in both Zld antibody replicates and the GFP antibody replicate. Comparison of Zld-binding sites between the embryo and neuroblasts were performed by intersecting high-confidence bound regions from the Zld antibody ChIP in the embryo and Zld antibody ChIP in the neuroblasts. Shared regions had 10bp overlap between the embryo and neuroblasts, and unique regions had less than 10bp overlap (Table S2). Visualization of genomic data was achieved by generation of z score-normalized bigWig files (Kent et al., 2010) from merged read coverage of replicates and displayed using Gviz (Hahne and Ivanek, 2016) and the UCSC Genome Browser (http://genome.ucsc.edu) (Kent et al., 2002; Raney et al., 2014). z score-normalized bigWigs were created by subtracting the mean read coverage (counts) from merged replicate read counts in 10bp bins across the entire genome and dividing by the standard deviation.

Heatmaps were generated using deepTools2 (Ramírez et al., 2016) with z score-normalized bigWig files. Average signal line plots were generated using seqplots from z score-normalized bigWig files at 10 base-pair resolution (Stempor and Ahringer, 2016). Genomic annotations were performed with the Bioconductor R package ChIPseeker (Bioconductor version 3.9, ChIPseeker version 1.18.0) using default settings and the BDGP dm6 genome through TxDb.Dmelanogaster.UCSC.dm6.ensGene package (BDGP version 1.4.1, TxDb version 3.4.4). TSS regions were redefined as −500bps to +150bps. Peak to nearest gene assignments were done using default settings of the annotatePeak() function.

To test for enrichment of motifs, *de novo* motif searches were done using the MEME-suite (version 5.1.1) (Bailey, 2011) and Hypergeometric Optimization of Motif Enrichment (HOMER, version 4.11) (Heinz et al., 2010). These programs identified motifs enriched in the input relative to shuffled or randomized regions of the genome. The *de novo* motifs were matched to known motifs from the JASPAR (http://jaspar.genereg.net) and DMMPMM (http://autosome.ru/DMMPMM) databases by the programs. Motifs are given a p-value or E-value (the p-value multiplied by the number of candidate motifs tested) indicating the confidence of the enrichment relative to the control sequences. Additional motif searches were done using the Biostrings package in R (version 2.50.2). The vcountPattern() function was used to tally the number of regions containing at least one occurrence of a motif and the vmatchPattern() was used to locate regions containing a motif.

To assess ChIP-seq and ATAC-seq replicate reproducibility, the number of reads overlapping each peak was quantified using featureCounts v1.6.4 (Liao et al., 2013). Log_2_(counts) overlapping each peak was plotted and the Pearson correlation calculated for each pair of replicates.

### Assay for Transposase-Accessible Chromatin

The protocol for ATAC-seq on larval brains was adapted from a protocol previously used for ATAC-seq on single embryos (Blythe and Wieschaus, 2016). *brat^11/Df(2L)Exel8040^* larvae were aged for 5-6 days at 25°C prior to dissecting brains in Schneider’s medium (Fisher, Hampton, NH, Cat # 21720001). 5 dissected brains were transferred to the detached cap of a microcentrifuge tube containing 10 μL cold lysis buffer (10 mM Tris, pH 7.5; 10 mM NaCl; 3 mM MgCl2; 0.1% NP-40). Under a dissecting microscope, brains were homogenized with the blunted tip of a pasteur pipette. The cap was transferred to a microcentrifuge tube containing 40 μL of additional lysis buffer. Tubes were spun down for 10 minutes at 500g at 4°C. Supernatant was removed under a dissecting microscope. This nuclear pellet was used to prepare ATAC-seq libraries using the Nextera DNA Library Preparation Kit (Illumina, San Diego, CA, Cat #FC1211030). The pellet was suspended in 5 μL buffer TD before adding 2.5 μL water and 2.5 μL tagment DNA enzyme. Samples were incubated at 37°C for 30 minutes. Tagmented DNA was purified using the Minelute Cleanup Kit (Qiagen, Hilden, DE, Cat #28004). DNA was amplified with 12 cycles of PCR using the NebNext Hi-Fi 2X PCR Master Mix (New England Biolabs Inc, Ipswich, MA, Cat #M0541S). Following PCR, DNA was purified using a 1.2X ratio of Axyprep magnetic beads (Axygen, Corning, NY, Cat #14223151). Library quality and tagmentation were assessed on an Agilent Bioanalyzer before pooling and submitting for 150 bp paired-end sequencing on a Illumina HiSeq 4000 at NovoGene.

Chromatin accessibility during the transition from type II neuroblast to INP was characterized using a temperature-sensitive system for synchronous type II neuroblast differentiation (Rives-Quinto *et al*. 2020). Brains were collected at 0, 6, 12, 18 or 24 hours following temperature shift, capturing intermediate stages during the transition from type II neuroblasts to INP. Following temperature shift, brains were collected and ATAC-seq libraries prepared as described above.

### ATAC-seq Analysis

Adapter sequences were trimmed from raw sequence reads using NGmerge (Gaspar, 2018). Reads were aligned to the *D. melanogaster* genome (version dm6) using Bowtie2 using the following parameters: --very-sensitive, --no-mixed, --no-discordant, -X 5000, -k 2 (Langmead and Salzberg, 2012). Aligned reads were filtered to include only reads with a mapping quality score > 30. Reads aligning to scaffolds or the mitochondrial genome were discarded. To identify fragments that likely originated from nucleosome-free regions, fragments were filtered to include only those < 100 bp, as previously described (Buenrostro et al., 2013). All downstream analysis and visualization was performed using these accessible fragments. To call peaks on accessible fragments, accessible fragments from both replicates were merged and MACS2 was used with the following parameters: -f BAMPE, --keep-dup all, -g 1.2e8, --call-summits. Deeptools was used to calculate genome coverage and generate bigWig files used for genome browser tracks and metaplots.

To compare Zld binding to chromatin accessibility in the embryo and neuroblasts, neuroblast-specific, embryo-specific and shared Zld peaks were merged to create a set of all Zld peaks detected in either cell type. The number of ChIP-seq or ATAC-seq reads overlapping each peak was quantified using featureCounts for the following datasets: neuroblast Zld ChIP, nuclear cycle 14 embryo Zld ChIP (Harrison et al., 2011), neuroblast ATAC and stage 5 embryo ATAC (Nevil et al., 2020). DESeq2 was used to compare ChIP-seq or ATAC-seq profiles between the embryo and neuroblast and identify differentially bound or differentially accessible regions between the two cell types (Love et al., 2014). Log_2_ fold-change values calculated by DESeq were used to correlate differences in binding with differences in accessibility between the two cell types.

## Data availability

Sequencing data have been deposited in GEO under accession code GSE150931.

## Acknowledgments

We thank the Bloomington Stock Center, and the Drosophila Genome Resource Center for providing reagents and fly lines. We acknowledge the University of Wisconsin-Madison Biotechnology Center and the NUSeq Core Facility for sequencing. Experiments were supported by grants from the National Institutes of Health: R01 NS111647 (M.M.H. and C.Y.L.), R35 GM136298 (M.M.H), and R01 NS107496 (C.Y.L) and a Vallee Scholar Award (M.M.H).

## Author Contributions

E.D.L., H.K., D.C.H., C.Y.L. and M.M.H conceived the study. E.D.L., H.K., T.J.G., C.M.O., D.C.H., and J.M.S. performed the experiments. E.D.L., H.K., T.J.G., C.M.O., D.C.H., J.M.S., C.Y.L. and M.M.H. analyzed the data. E.D.L., C.Y.L. and M.M.H wrote the manuscript. E.D.L., H.K., C.Y.L. and M.M.H. revised the manuscript.

