## Supplemental Figures for "Cell-type-specific chromatin occupancy by the pioneer factor Zelda drives key developmental transitions in *Drosophila*"

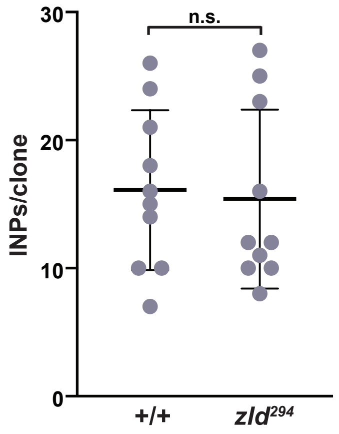


**Figure S1 - related to Figure 1: INP number in *zld*-null mutant neuroblast clones is indistinguishable from wild-type.** Number of INPs in either wild-type or *zld*$^{294}$ type II neuroblast clones as marked by Dpn and Ase expression (Dpn+Ase+) . +/+ = 16.1 ± 6.2 INPs and *zld*$^{294}$ = 15.4 ± 7 INPs; n = 10 clones per genotype. Comparison done using a two-tailed Student's t test; n.s. = p-value >0.05.

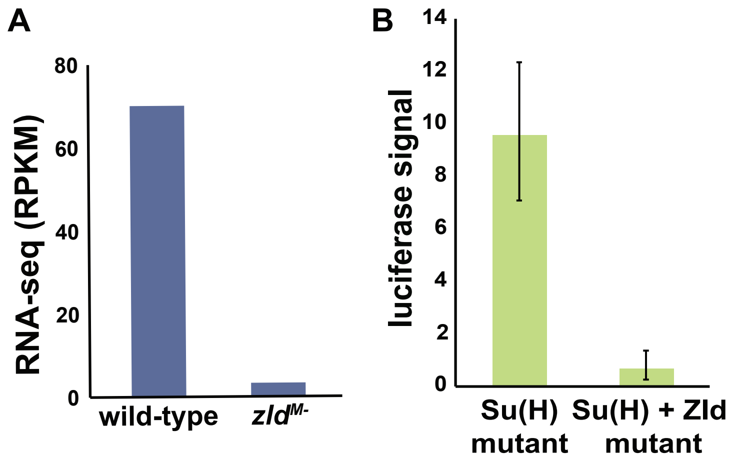


**Figure S2 - related to Figure 2: *dpn* is a Zld-target gene in the early embryo.** A. *dpn* mRNA levels decrease in stage 5 embryos lacking maternally deposited *zld* (*zld*$^{M-}$) compared to wild-type. (Schulz et al. 2015). B. Luciferase expression of the *dpn* GFP:luciferase reporters in 2-3 hr AEL (after egg laying) embryos require Zld-binding sites for expression. Error bars represent the standard deviation.

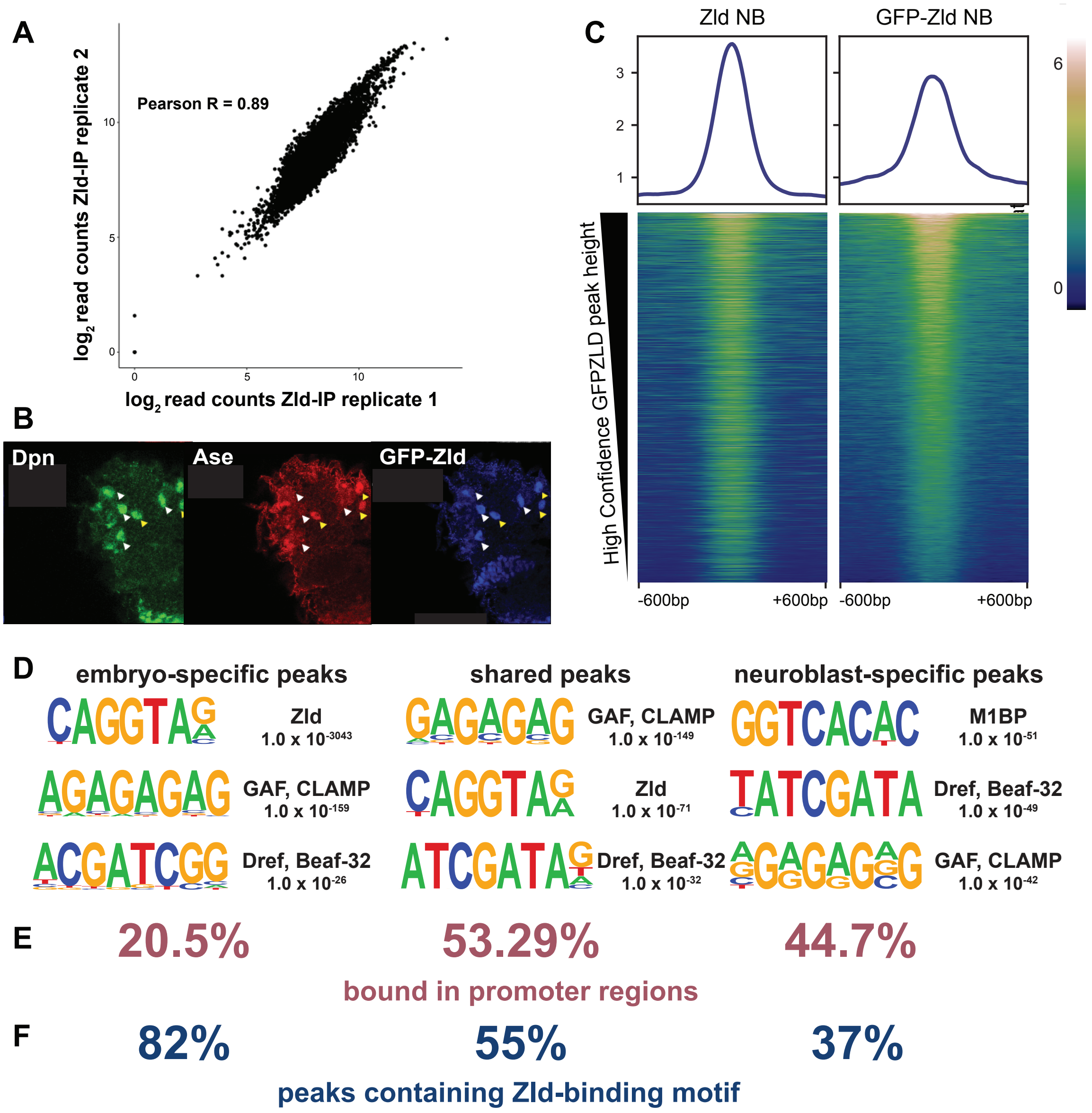


**Figure S3 - related to Figure 3-4: ChIP-seq for GFP-Zld confirms neuroblast-specific Zld-bound regions.** A. Pearson correlation plot of read coverage for the two Zld-antibody replicates for ChIP-seq from *brat* mutant brains shows high correlation between replicates. B. Third instar larval brain stained for Ase, Dpn and GFP-Zld. GFP-Zld is expressed in type I neuroblasts (yellow arrowhead) and type II neuroblasts (white arrowhead). C. Heat maps centered on the ChIP peak with 600bp flanking sequence for peaks bound by the anti-Zld antibody (Zld NB) and the anti-GFP antibody (GFP-Zld NB). Colors indicate relative ChIP z score. Average z score is shown above each heat map. The highest bound regions identified by the Zld antibody are also the highest bound regions identified by the GFP antibody. D. Motif enrichment for the high-confidence embryo-specific peaks, shared peaks and neuroblast-specific peaks identified by both Zld and GFP antibodies as determined by HOMER. The same motifs are enriched as when only regions bound by the Zld antibody are analyzed. E. Fraction of high-confidence Zld-bound regions (identified using anti-Zld and anti-GFP antibody) bound in promoters (-500bps to +150bps). There is an enrichment for Zld binding to promoters in the regions specific to the neuroblasts and shared regions between the embryo and neuroblasts. F. Percent of peaks in each class that contain a variant of the Zld-binding motif (NAGGYAN). The Zld-binding motif is more enriched in regions specific to the embryo and shared regions.

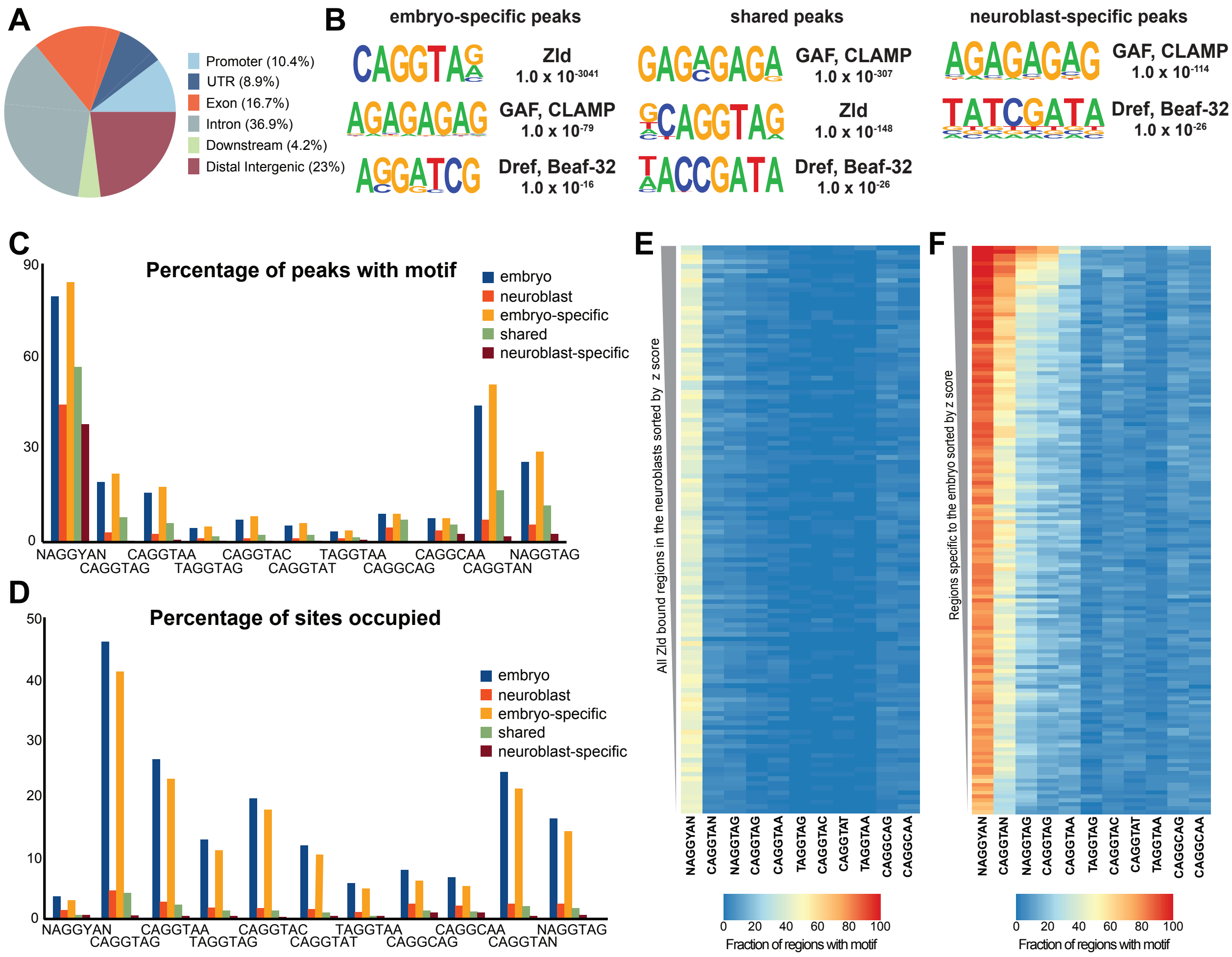


**Figure S4 - related to Figure 4: The canonical Zld-binding motif is not enriched in regions specifically bound by Zld in the neuroblasts.** A. Genomic distribution of random 200bp regions of the *Drosophila* genome (shuffled unique neuroblast Zld-bound regions). Promoter regions defined as -500bps to +150bps from the TSS. B. Motif enrichment for the high-confidence embryo-specific peaks, shared peaks and neuroblast-specific peaks using the anti-Zld antibody as determined by HOMER. The canonical Zld-binding motif is not enriched in regions specific to Zld binding in the neuroblasts. C. Percent of peaks containing the Zld-binding motif variant indicated for each class of Zld-bound peaks. D. Percent of motif variants in the genome occupied in each class of Zld-bound regions. E. Enrichment of Zld-binding motif variants in all Zld-bound regions identified in type II neuroblasts. Peaks are sorted from highest (top) to lowest (bottom) z score and binned into groups of 100. Color of the cell represents the percent of peaks in each bin containing the indicated motif within the 200bp peak region. There is a low level of enrichment of all specific variants on the Zld-binding motif at regions bound by Zld in the neuroblasts. The most enriched variant is a degenerate motif encompassing all variations on the Zld-binding motif (NAGGYAN). F. Enrichment of Zld-binding motif variants in all Zld-bound regions specific to the early embryo. Peaks are sorted from highest (top) to lowest (bottom) z score and binned into groups of 100. Color of the cell represents the percent of peaks in each bin containing the indicated motif within the 200bp peak region. While the degenerate motif (NAGGYAN) is the most enriched motif of regions specific to Zld binding in the embryo, the specific variant with the most enrichment among embryo-specific regions is CAGGTAN.

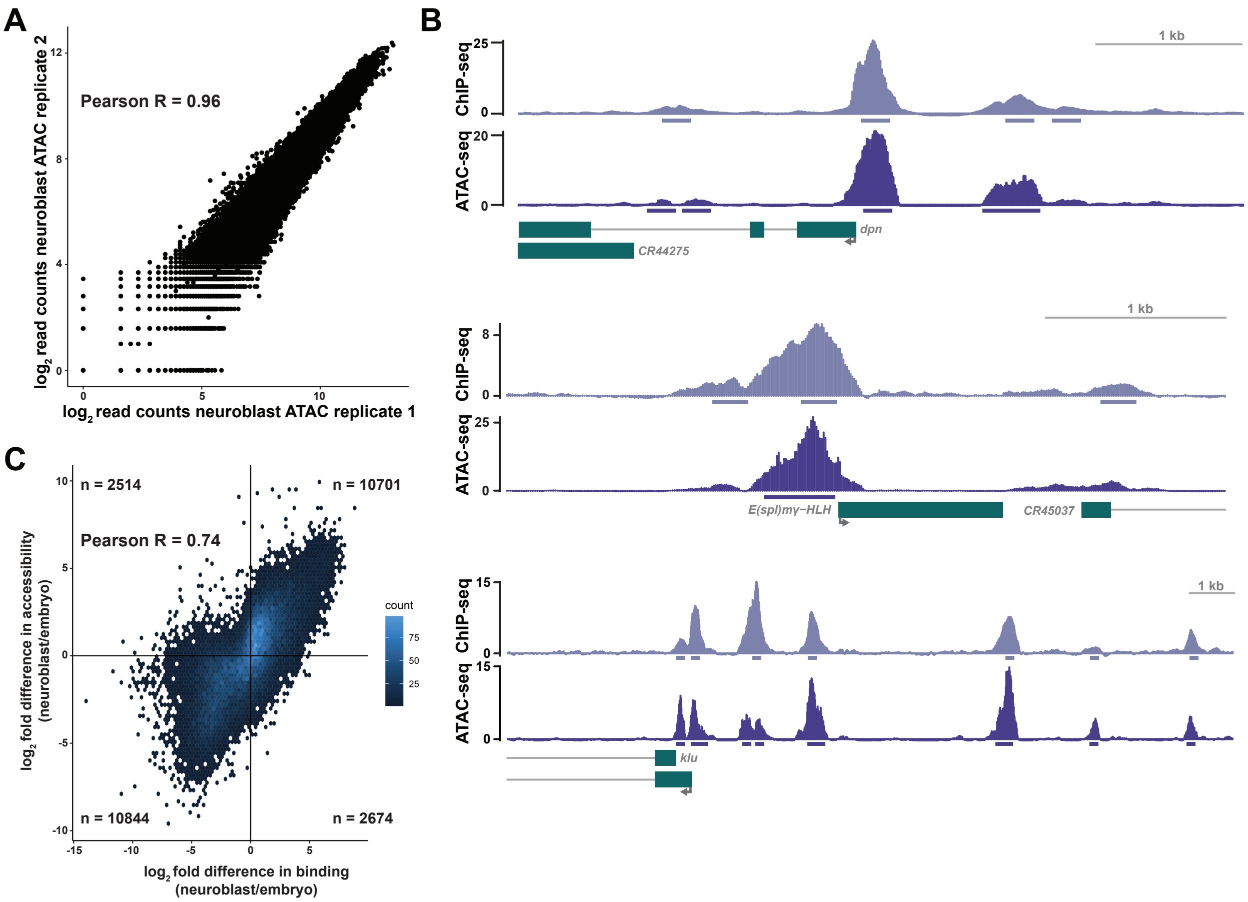


**Figure S5 - related to Figure 5: Zld-binding correlates with chromatin accessibility in the embryo and type II neuroblasts.** A. Pearson correlation plot of read coverage for ATAC-seq replicates on *brat* mutant brains on a $\log_2$ scale shows high correlation between replicates. B. Genome browser tracks of Zld-binding (ChIP-seq) and chromatin accessibility (ATAC-seq) from type II neuroblasts at the *dpn, E(spl)mγ-HLH* and *klu* loci. Peak regions are shown below the tracks. C. Differences between Zld binding (ChIP-seq) in the embryo and type II neuroblasts correlate with differences in chromatin accessibility. $\log_2$ fold difference in Zld binding on the x-axis compared to $\log_2$ fold difference in chromatin accessibility on the y-axis. Color represents relative count.

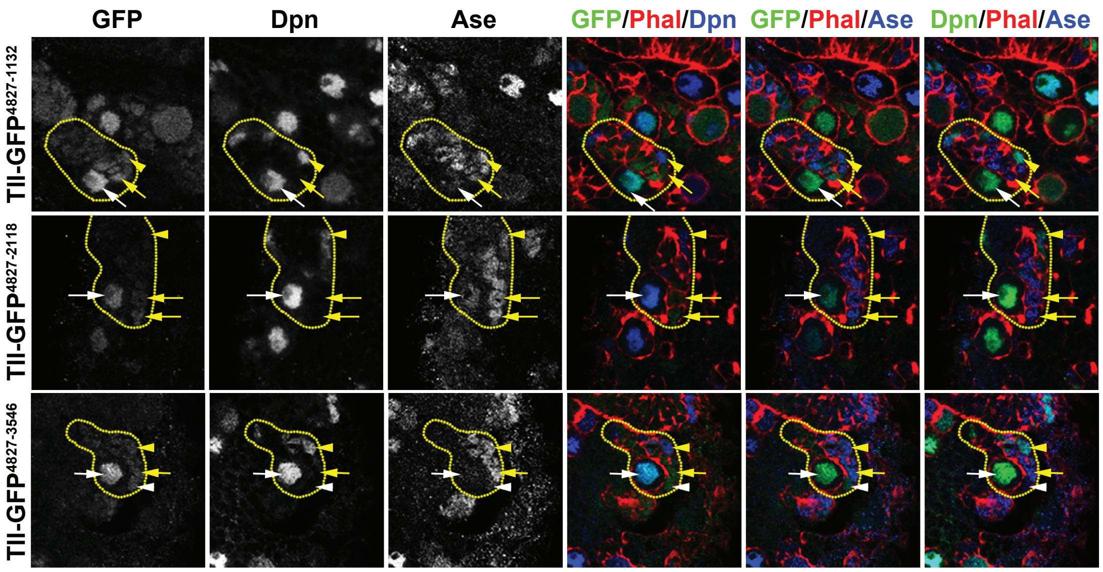


**Figure S6 - related to Figure 6: TII neuroblast-specific Zld bound enhancer drives reporter expression in neuroblasts.** Representative images of GFP expression in type II neuroblasts of animals expressing transgenes containing truncations of the upstream regulatory region of *tll* and DSCP driving a GFP reporter. Staining for markers of neuroblasts are also shown. Type II neuroblast lineages are outlined with a dashed yellow line. Yellow arrowheads represent INPs, white arrowheads represent Ase- immature INP, yellow arrows represent Ase+ immature INP and white arrows represent type II neuroblasts.

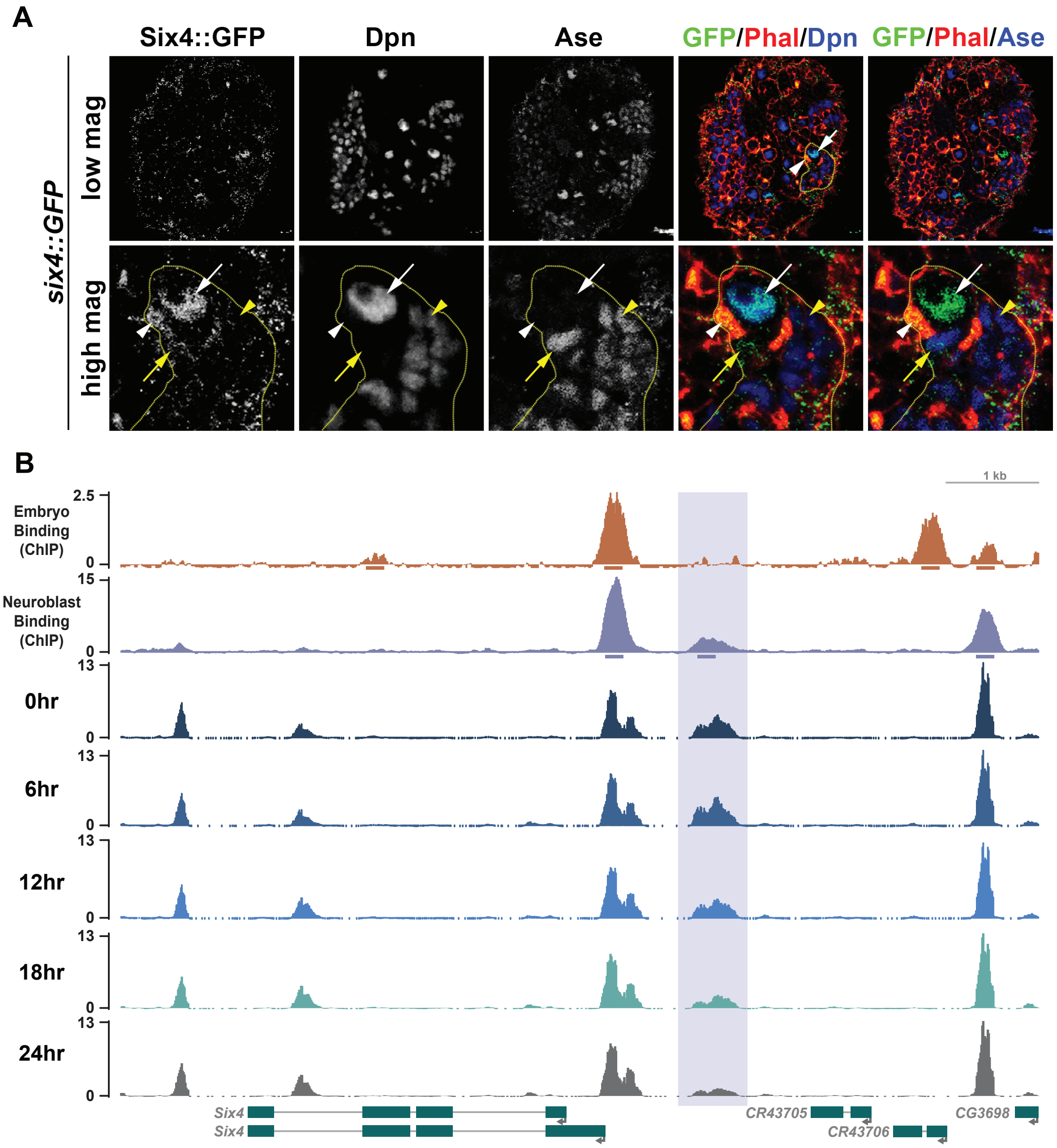


**Figure S7 - related to Figure 6: Six4 expression is limited to the neuroblasts and a neuroblast-specific Zld bound *Six4* enhancer loses accesibility throughout differentiation.** A. Representative images of GFP expression in type II neuroblasts of animals expressing transgenes containing a *Six4::GFP* transgene at low (top) and high (bottom) magnification. Staining for markers of neuroblasts are also shown. Type II neuroblast lineages are outlined with a dashed yellow line. Yellow arrowheads represent INPs, white arrowheads represent Ase- immature INP, yellow arrows represent Ase+ immature INP and white arrows represent type II neuroblasts. B. Genome browser tracks of Zld ChIP-seq from the early embryo and type II neuroblasts and ATAC-seq of the *Six4* locus from *brat* mutant brains at the indicated time points following a temperature shift that initiates synchronous differentiation. 200 bp regions surrounding the ChIP peak summits are shown below the top two tracks.
